# Lineage memory shapes viral resistance barriers in human skin

**DOI:** 10.1101/2025.06.27.661951

**Authors:** Laura C. Van Eyndhoven, Grant Kinsler, Jingchao Zhang, Aoife O’Farrell, Rim Abderrahim, Apurv Srivastav, Catherine G. Triandafillou, Todd M. Greco, Kenneth S. Zaret, Ileana M. Cristea, Megan H. Orzalli, Nir Drayman, Abhyudai Singh, Arjun Raj

**Affiliations:** Department of Bioengineering, School of Engineering and Applied Sciences, University of Pennsylvania, Philadelphia, PA, USA; Institute for Regenerative Medicine, Penn Epigenetics Institute, Department of Cell and Developmental Biology, Perelman School of Medicine, University of Pennsylvania, Philadelphia, PA, USA; Department of Molecular Biology, Princeton University, Princeton, New Jersey, USA; Department of Electrical and Computer Engineering, University of Delaware, Newark, Delaware, USA; Department of Medicine, Division of Infectious Diseases and Immunology, University of Massachusetts Chan Medical School, Worcester, MA, USA; Program in Innate Immunity, University of Massachusetts Chan Medical School, Worcester, MA, USA; Department of Molecular Biology and Biochemistry, University of California, Irvine, Irvine, CA, USA; Department of Genetics, Perelman School of Medicine, University of Pennsylvania, Philadelphia, PA, USA

## Abstract

Individual cells within a given population exhibit striking variability in viral susceptibility, but it remains unknown whether this heterogeneity reflects memories encoded into the cellular lineage or true probabilistic variability. We used multi-color lineage tracing in a human primary organotypic skin model to reveal that viral resistance is encoded within specific cellular lineages. These lineages create distinct boundaries that block viral spread. Our lineage analyses *in vitro* confirmed that viral susceptibility exhibits strong heritability across cell generations, with siblings and cousins displaying remarkably similar infection outcomes. ATAC and proteomics profiling of resistant and susceptible clones revealed distinct epigenomic and proteomic states, with the transcription factor AP-1 emerging as a potential central regulator of lineage-encoded viral resistance. Inducing AP-1 activity with PMA rendered cells resistant to viral infection, suggesting a causative role in mediating resistance memory. Our findings demonstrate that antiviral resistance in human skin cells is encoded within cellular lineages and preserved through cell divisions, revealing how cell memory may shape infection dynamics and viral containment in tissues.

## Introduction

Heterogeneity in cellular responses to challenges is a common feature of biological systems. The innate immune system is a salient example, with individual cells showing marked variation in their expression of key factors. There is a growing appreciation that cellular lineage can explain seemingly probabilistic cellular variability and responses in areas like oncogenesis and therapy resistance in cancer [1–3], hematopoiesis [4,5], and innate immune signaling [6,7], reflecting the encoding of cellular “memory”. However, little is known about whether lineage memory can affect the organization of innate immune responses at the tissue level.

Numerous studies have documented substantial variability in the expression of key antiviral genes when host cells are exposed to viral particles [8,9]. This intrinsic heterogeneity is often amplified through positive feedback mechanisms, resulting in highly skewed or bimodal expression patterns that suggest the involvement of complex regulatory switches and epigenetic modifications [6,10]. Recent studies using single-cell cloning and retrospective clone tracing have begun to reveal that certain cellular states confer reduced infectability [11,12], with these states exhibiting varying degrees of memory across cell divisions. However, the extent to which this memory operates within complex tissue environments, where cell-cell interactions and spatial organization play critical roles, and how it relates to the observed heterogeneity in antiviral factor expression, remain important unresolved questions.

A key organizational principle for cellular heterogeneity that is relevant to innate immune responses is the timescale over which cells fluctuate between different states. Cell memory operates across a spectrum, from rapidly changing states that persist for only hours to stable epigenetic configurations that persist through multiple cell generations. Rapid fluctuations are often considered stochastic noise, but longer-lasting fluctuations represent a form of cellular memory, as cells within a lineage will share these states and thus potentially exhibit correlated behaviors [1]. The timescale of fluctuations can have a strong effect on how tissue responses are spatially organized during viral infection. Short-lived memory would generate a “salt and pepper” pattern of cellular variability throughout the tissue, with resistant and susceptible cells intermingled randomly. An example could be the expression of interferon and interferon-stimulated-genes, which are known to fluctuate rapidly [1,7], and may contribute to an overall anti-viral activation across the whole tissue via secretion and diffusion. In contrast, longer-lasting memory gives rise to cell lineages that share similar phenotypes. Memory formation has emerged as an important aspect of innate immunity, for instance, in trained immunity, where an initial stimulus can lead to enhanced responsiveness to subsequent stimuli [13]. In tissue contexts, tissue-resident stem cells and their progeny may encode epigenetic states that locally manifest as coherent patches of cells exhibiting similar phenotypes. If one of these clonal patches were resistant to infection, it might establish defined barriers that could restrict viral spread. However, without methodologies to trace cellular lineages with single cell resolution, it remains challenging to determine the timescales of these fluctuating responses and their functional consequences, particularly in complex *in vivo* environments.

Here, we developed a multi-color, genetically-encoded, fluorescence-based lineage tracing approach to directly address whether viral resistance in human skin is governed by cellular memory and at what timescales this memory operates. We used single-molecule RNA fluorescence *in situ* hybridization (smRNA FISH) in human organotypic skin constructs challenged with viruses to demonstrate that infection dynamics are dictated by slow-fluctuating cell states that maintain viral susceptibility or resistance across multiple generations. Consequently, these lineages create distinct boundaries that block viral spread. Through quantitative analysis of sibling and cousin cells, we determined that this resistance memory has a half-life of approximately 35 cell divisions—establishing a mechanism for how clonal patches can form effective barriers against viral spread in tissues. Multi-omic profiling of resistant and susceptible clones revealed that this memory is associated with epigenetic signatures, with the AP-1 transcription factor emerging as a central regulator of viral resistance memory. Proteomic analysis corroborated the ATAC-seq findings, uncovering widespread differences in antiviral immune factors between resistant and nonresistant clones. Our findings support the possibility that viral spread in human skin is controlled by spatial organization resulting from lineage-encoded antiviral memory rather than rapid cellular fluctuations and communication.

## Results

### Organotypic skin cultures allows for studying lineage-encoded memory in tissues

Organotypic skin cultures are an elegant model in which to determine whether lineage-encoded memory affects viral infection in a tissue context (Figure 1A). As occurs *in vivo*, stem cells reside at the base, producing lineages of cells that grow upward through the tissue, with individual lineages remaining spatially segregated [14–16]. Moreover, organotypics provide the flexibility to incorporate genetically modified cells (e.g., lineage-labeled cells) while maintaining complex tissue architecture [17,18]. We generated organotypic skin cultures by seeding human primary keratinocytes on devitalized dermis obtained from healthy donors [16]. Upon seeding, the keratinocytes self-organize by forming a distinct proliferative stem cell zone on top of the basement membrane (Figure 1B). Cultured in an air-liquid interface, the cells take about two weeks to make up a 3D tissue architecture that recapitulates human skin morphology and function [16,17]. The outermost layer of the epidermis is composed of terminally differentiated, non-nucleated keratinocytes, which form a robust barrier that prevents viral entry and replication. Since viruses require nucleated host cells to replicate, we introduced viruses directly to the nucleated cells at the base by puncturing the surface with a microneedling device, thereby bypassing the non-permissive outer layer, as described earlier [14].

**Figure 1:**
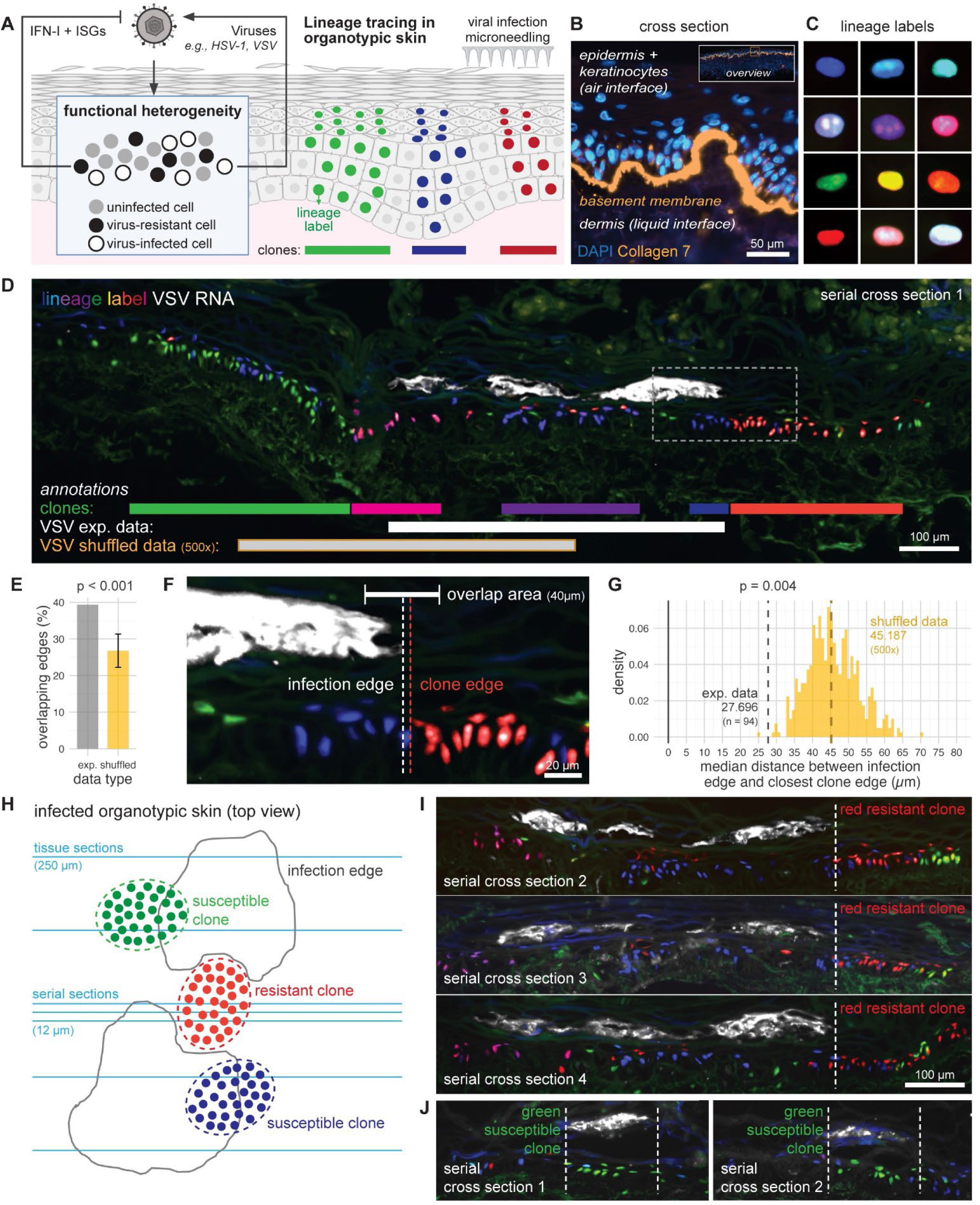
Antiviral cell states show strong memory across clonal lineages. A) : Schematic of lineage tracing in organotypic skin challenged with live replicating viruses. Lineages are traced using fluorescent nuclear lineage labels. Viral infection is established upon microneedling. B) : Cross section of organotypic skin with immunofluorescence staining for collagen 7 to visualize the basement membrane. C) : Examples of the nuclear fluorescent lineage labels. D) : Representative example of rVSV-M51R-infected organotypic cross section (serial section 1). Clones were traced using the lineage labels. Viral replication was quantified using smRNA FISH. Annotations include the clones and VSV locations in the horizontal plane. The experimental (abbreviated with exp.) data for the VSV locations match the actual location of the VSV replication sites, whereas the shuffled data was computed randomly 500 times. E) : Quantification of the overlapping edges (%) for the experimental data (n = 3) and shuffled data. F) : Zoom in on the example cross section (dashed box in panel D), indicating the infection edge, clone edge, and overlap area of 40 µm. G) : Quantification of the median distance between infection edge and closest clone edge for the experimental data and the 500 randomly computed (shuffled) data. The p-value represents the empirical p-value derived from comparing to shuffled data. H) : Schematic of infected organotypic skin (top view) with infected regions indicated by the gray infection edges. Clones are represented by the different color dots, of which one example edge is indicated with the blue dotted line. The light blue horizontal lines represent the tissue sections, which are either 250 µm apart, or only 12 µm in case of the serial sections. I) : Continuation of the serial sections (2-4), with VSV infection edge annotated, highlighting a red resistant clone. J) : Second representative example on lineage-encoded memory across 2 serial sections, highlighting a green susceptible clone.

To determine the extent of lineage-encoded memory in this system, we devised a genetically-encoded nuclear fluorescence cell lineage labeling strategy to track cell lineages across generations. This strategy leverages the expression of varying combinations of three fluorescent proteins that localize exclusively in the nucleus (Figure 1C). By restricting the expression of fluorescent proteins to the nucleus, the cytoplasm remained optically accessible for additional fluorescent quantifications, such as fluorescent viral reporter proteins and gene expression using smRNA FISH. This setup allowed for seamless integration with other fluorescence-based analyses, including quantification of viral replication in organotypic skin.

### Antiviral cell states show strong memory across clonal lineages

We first tested whether we could observe clone-to-clone variability in virus susceptibility in tissues using our nuclear fluorescence cell lineage labeling strategy. We infected human organotypic skin with recombinant vesicular stomatitis virus VSV carrying the M51R point mutation (rVSV-M51R, later referred to as VSV). This mutation results in a delayed ability to inhibit host gene expression, as well as an inability to suppress type I interferon (IFN-I) gene expression [19,20]. As a result, this virus not only replicates efficiently but also permits a more relevant interrogation of host antiviral defense mechanisms, enabling us to study both viral spread and the spatial organization of innate immune responses in tissue. We quantified viral replication using smRNA FISH targeting the L viral transcript after 3 days of infection (see Materials and Methods). We tracked clones using the nuclear fluorescent lineage labels. Towards the stratum corneum (the outer layer of the skin), cells start to lose their nuclei, and subsequently their fluorescent lineage labels. Stem cells give rise to differentiated cells via a vertical expansion from the base [21]; hence, we annotated the clonal lineages in the stem cells at the bottom of the layer with the assumption that the cells above them were of the same lineage.

Notably, VSV infection was largely restricted to differentiated cells in the upper layers, while basal stem cells often remained uninfected (Figure 1D). This pattern is consistent with previous findings that stem cells express higher levels of antiviral factors, providing them with intrinsic protection [22]. As a result, viral replication occurred primarily in differentiated progeny, allowing us to assess how lineage-derived properties influence susceptibility to infection. Visually, the borders of viral replication sites often coincided with clonal boundaries, suggesting that certain clones consisted predominantly of virus-resistant cells (Figure 1D). To quantify these interactions, we assessed how often boundaries of viral replication coincided with clonal boundaries (Figure 1E). If viral replication would not be governed by lineage-encoded memory, VSV replication would occur in the skin cells independently of the location of clonal lineages; in that case, their edges would not overlap any more than expected by random chance. To test that, we analyzed 26 organotypic skin cross-sections, obtained from three independent biological replicates, containing 47 VSV infection sites and 240 clones. 39.4% of all VSV edges overlapped with the edge of its nearest clone, compared to 26.8% of the computationally shuffled data (p < 0.001, see Methods for details on shuffling analysis). We counted edges as overlapping when they were less than 20 µm apart (Figure 1F). Shuffled data was obtained by computationally shuffling the VSV annotations 500 times at random. Note that we only expected a small subset of clones to be resistant to infection; hence, the enrichment of overlapping edges from 26.8% to 39.4% was in line with our expectations. We also measured the median distance between the edge of infection and the closest clone edge to be 27.7 µm, compared to 45.2 for the shuffled data (p = 0.004, Figure 1G).

Next, we analyzed data from serial sections to further support our hypothesis of lineage memory governing viral replication spread (Figure 1H). We reasoned that if all the cells in a clone were resistant, the boundary between the edge of VSV infection and the putatively resistant clone would overlap across multiple, if not all, serial sections. We found numerous examples of this behavior (as well as examples of clones that were not resistant, even fully susceptible), with two representative examples highlighted. The first example shows a red resistant clone in multiple serial sections (Figure 1I). The second example shows a green susceptible clone in multiple serial sections (Figure 1J). Overall, we conclude that viral replication in tissues is governed by lineage-encoded memory, resulting in clear boundaries between infected and uninfected regions.

### Antiviral lineage-encoded memory manifests as slow-fluctuating cell states

To quantify the time scale over which the memory of relevant cell states persisted, we turned to *in vitro* models of infection combined with lineage tracing and single-molecule RNA FISH readouts. VSV infection and replication *in vitro* can occur rapidly (Supplementary Figure 1A). At 1 hour post-infection with rVSV-M51R (later referred to as VSV), at an MOI of 2, individual viral RNA spots could be detected. At 6 hours post-infection, a small fraction of cells had over 600 VSV mRNA counts (Supplementary Figure 1B). After 24 hours, roughly half of the cells experienced high viral loads, while the remaining fraction showed barely any viral replication despite being infected (Supplementary Figure 1C).

Next, we traced clones using the lineage labels in virus-infected cells *in vitro* in the same manner as we did in the organotypic skin. Tracing large clones (sometimes over 20 cells per clone per cross-section) in organotypic skin was relatively easy because of their confinement by the tissue architecture. However, *in vitro*, cells could move around substantially over the course of generations. The combination of a lacking spatial confinement and a limited color diversity of lineage labels increases the chance of unrelated cells of a similar color being in close proximity to each other, especially at high generation numbers. We therefore opted to only allow cells to undergo 1 or 2 divisions *in vitro*, leading to a distribution of cells per clone of 2 to 4 on average (Figure 2A). Next, we calculated the coefficient of variance (CV) of VSV mRNA counts as a measure of variability in viral replication within cells from the same lineage. A small CV would indicate that cells within a clone had relatively similar levels of infection, whereas a large CV would indicate more variance in those levels. In line with the results obtained in organotypic skin, we found smaller CVs within clones compared to scrambled data (Figure 2B). Of note, while it is possible for cells to divide during infection, VSV replicates extremely rapidly, reaching high levels of viral mRNA within just 6 hours. In contrast, cell division occurs over much longer timescales (∼24 hours), making it unlikely that the observed similarity in viral loads within clones is primarily due to inheritance of viral load from an already-infected parent cell. We thus conclude that lineage-encoded memory governs viral replication, resulting in cells within a clone to experience similar quantities of viral replication.

**Figure 2:**
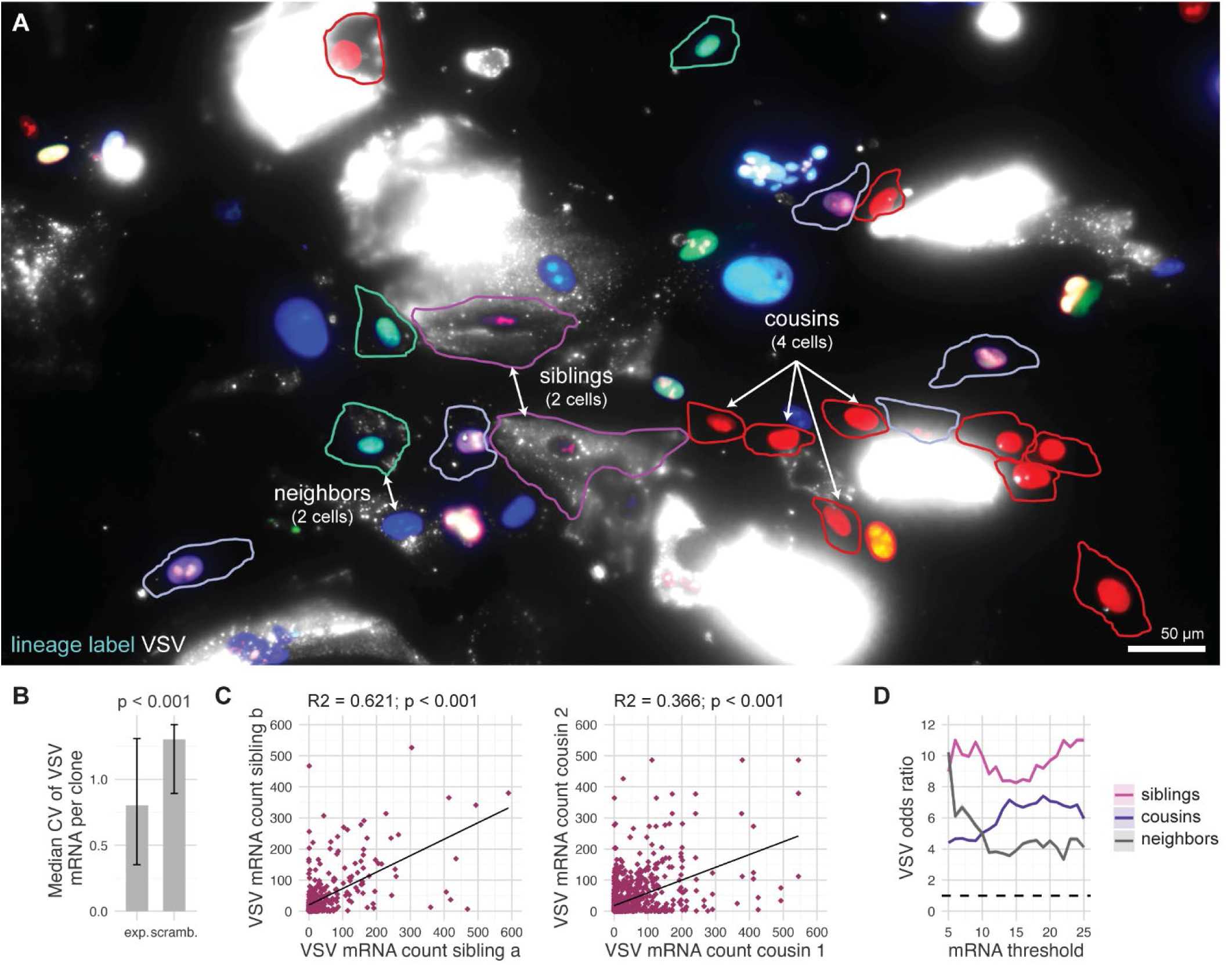
Lineage-encoded memory of viral susceptibility is a reflection of slow-fluctuating cell states. A) : Representative microscopy image of human primary fibroblasts infected with rVSV-M51R for 24 hours. Cell lineages are traced using nuclear fluorescent lineage labels. Viral replication is quantified using smRNA FISH. Annotations highlight examples of siblings, cousins, and neighbors. B) : Bar graphs representing the median and interquartile range of coefficient of variance (CV) of VSV mRNA per clone from the experimental (exp.) data, compared to the scrambled (scramb.) data (n = 2). C) : Correlations plots of the VSV mRNA counts comparing siblings and cousins. D) : Odds ratios for shared VSV mRNA expression among cell pairs (siblings, cousins, neighbors) across varying expression thresholds.

Additionally, we compared levels of viral replication between siblings and cousin cells, which we defined based on shared lineage labels: sibling cells arose from the same most recent division event (i.e., daughter cells of the same mother), while cousin cells shared a common ancestor but were separated by at least one additional division. VSV mRNA counts correlated in both siblings and cousins (p < 0.001) (Figure 2C). We next binarized the data and calculated the odds ratio for siblings, cousins, and neighbors (cells that are adjacent but lineage-unrelated) to compare the observed frequency of concordant states to that expected if viral mRNA expression states were independent within cell pairs. For a wide range of thresholds, the odds ratio for siblings and cousins was far above 1 (e.g., with values of 8 and 7, respectively, for a threshold of 15 VSV mRNA) (Figure 2D), again supporting that viral resistance and susceptibility is governed by a strong memory amongst the cells within a clone.

The odds ratio for neighboring cells being greater than 1 indicates a minor spatial correlation in infection patterns. Specifically, cells adjacent to highly infected cells are more likely to exhibit high viral loads themselves, suggesting local biases in viral spread between neighboring cells. However, this effect was small compared to the associations between siblings and cousins.

Next, to get a sense of the duration of memory, we modeled the memory as a two-state stochastic switching model in which cells can transition between an ON (susceptible to viral infection, corresponding with viral transcription) and an OFF (protected from viral infection, corresponding with no to low viral transcription) state. This model has two key parameters: the k_on_ and k_off_ rates, which serve as metrics quantifying the timescales at which the states fluctuate (see Methods for additional details). Our modeling calculated k_on_ and k_off_ values of roughly 0.02 day^−1^, suggesting that viral replication is governed by clone-to-clone heterogeneity encoded by slow-fluctuating cell states (Supplementary Figure 2A, B). In fact, at rates of 0.02 day^−1^, it takes approximately 35 days for 50% of the population to switch fate. Given that the cells roughly divide once a day, we can infer that the memory half-life is approximately 35 cellular divisions.

One natural explanation for the antiviral propensities of certain clones is that these clones may have activated known antiviral pathways. In that scenario, clones with increased antiviral gene expression would experience decreased levels of viral replication. IFN-Is are arguably the most potent antiviral cytokines, known for their strong antiviral effects [23,24]. IFN-I production is known to be highly heterogeneous, with some cells producing massive amounts upon activation and others producing none, as we observed across multiple cell types (Supplementary Figure 3A). However, a similar sibling and cousin cell analysis of heterogeneity in IFN-I (i.e., IFNβ) expression levels upon PolyIC activation revealed much lower sibling and cousin correlations, lower odds ratios, and higher k_on_ and k_off_ rates than viral RNA expression levels (Supplementary Figure 3B-E). Similar results for IFN-I expression correlations were obtained in cells infected with VSV (Supplementary Figure 4A-D). These analyses all pointed to much faster state transitions between IFN-I-high vs. IFN-I-low cells, showing that memory of IFN-I can only partially explain the apparently clonal memory of susceptibility to viral replication. Of note, we quantified a non-significant anti-correlation of IFN-I mRNA counts and VSV mRNA counts (R^2^ = −0.05, p = 0.052), suggesting that IFN-I production does not strongly contribute to functional antiviral protection in this context (Supplementary Figure 4E).

Given the limited connection between IFN-I memory and viral susceptibility, we next asked whether the activation of downstream antiviral pathways, quantified by MX1 smRNA FISH, might instead align with the boundaries of susceptibility. We found that uninfected cells near infected regions had much higher MX1 mRNA counts compared to uninfected cells near uninfected regions (median of 35 versus 2, respectively, n = 3; p < 0.001) (Supplementary Figure 5A-C). However, this response appeared independent of clone boundaries, indicating that antiviral pathway activation was spatially induced but not clonally restricted.

In conclusion, the strong similarities in viral load among clones and high variability between clones showed that viral susceptibility is dictated by slow-fluctuating cell states. Since antiviral responses, quantified by IFN-I production, were fast-fluctuating, we concluded that viral replication is primarily dictated by a cell state that is independent of the canonical antiviral activity of IFN-I production and subsequent ISG upregulation upon infection.

### Viral resistance displays strong memory in clones

To determine the molecular basis for the slow fluctuations that lead to clonal memory of viral susceptibility, we needed to isolate larger numbers of cells from a particular clone. We thus grew clonal populations arising from single cells. The first step was to test for viral susceptibility of these clones. To remove potentially confounding effects of IFN-I production that might affect viral replication dynamics within and between cells, we used herpes simplex virus 1 (HSV-1) infections for determining viral susceptibility. HSV-1 is a skin-tropic virus known to prevent IFN-I expression and signaling by its host (reviewed in [25]).

We generated single-cell clones of human primary skin fibroblasts by limiting dilution in 96 well plates (Figure 3A). To quantify viral replication, we used wildtype HSV-1 (strain 17) expressing YFP-ICP4. ICP4 is an immediate-early protein that is expressed as soon as the viral genome enters the nucleus [26]. To track viral susceptibility, replication, and disease progression during the first 24 hours of infection, individual clones were infected with this modified HSV-1 at a low multiplicity of infection (MOI) of 0.01-0.001. We observed two distinct outcomes at the single-cell level. In some cases, the fluorescent reporter signal remained confined to the nucleus, indicating the virus successfully entered the cell but failed to replicate and spread to neighboring cells [26]. In other cases, fluorescence extended beyond the nucleus into the cytoplasm, consistent with active viral replication and secondary infection of adjacent cells. We defined the former as *infection sites* (initially infected cells with nuclear fluorescence only) and the latter as *plaques* (clusters of infected cells resulting from successful viral spread).

**Figure 3:**
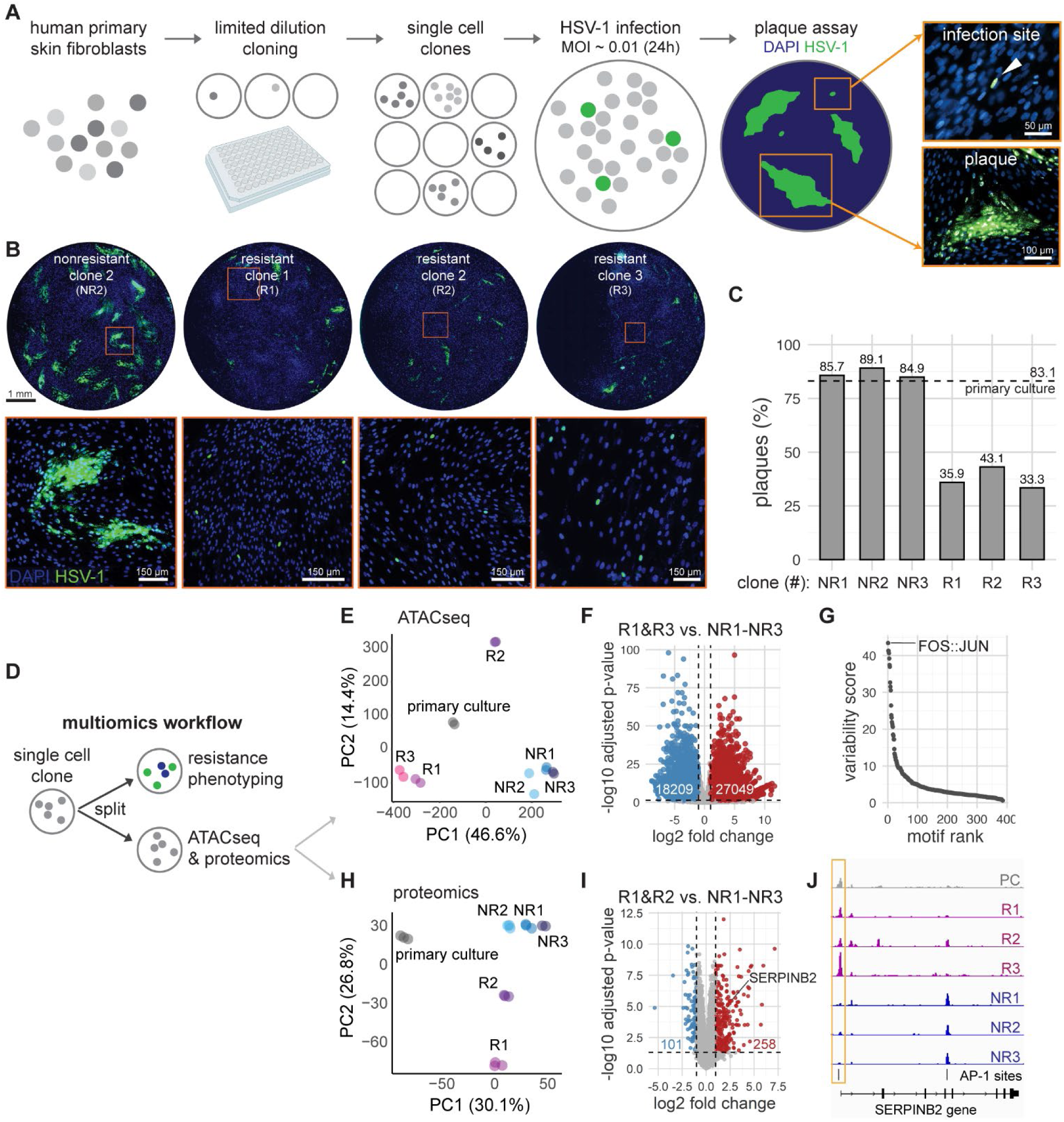
Clonal lineages show remarkable differences in viral resistance. A) : Schematic on the experimental approach for making single-cell clones and infecting them with HSV-1 ICP4-YFP to assess resistance. Once single virus particles enter the monolayer, after 24 hours, they either get restricted to their initial infected host cell, leading to a so-called infection site, or they continue to infect neighboring cells, resulting in the formation of a so-called plaque. B) : Example images of a nonresistant clone (NR2) and 3 resistant clones (R1, R2, R3), with their corresponding zoom in (orange boxes) to represent the plaque formation in the nonresistant clone and the infection sites in the resistant clones. Clones were infected with HSV-1 for 24 hours at an MOI of 0.01. C) : Bar graph of mean + standard deviation of plaque percentage per nonresistant and resistant clone. The dashed line represents the percentage of plaques obtained from the primary culture. D) : Schematic on the multi-omics workflow. E) : Dimensionality reduction analysis of the ATAC sequencing data on binned, normalized reads. F) : Volcano plots of the differential peaks comparing R1&R3 (2 resistant clones) versus NR1-NR3 (3 nonresistant clones). Red dots represent peaks that are differential in resistant clones, blue dots represent peaks that are differential in nonresistant clones. Thresholds: adj.P.Val < 0.05 & logFC > 1, −1. G) : Variability scores of the ChromVAR analysis, ranked from highest to lowest. H) : Dimensionality reduction analysis of the proteomics data on protein abundance profiles. I) : Volcano plots of the differential protein abundance comparing R1&R2 (2 resistant clones) versus NR1-NR3 (3 nonresistant clones). Red dots represent proteins that are differential in resistant clones, blue dots represent proteins that are differential in nonresistant clones. SERPINB2 is highlighted. Thresholds: adj.P.Val < 0.05 & logFC > 1, −1. J) : Representative ATAC sequencing tracks of the SERPINB2 locus. AP-1 binding site is depicted in the bottom row.

### Clonal variability in viral resistance is defined by an incomplete viral life cycle, despite equal susceptibility to entry

Upon infection, clones exhibited striking differences in their ability to resist viral spread (Figure 3B). Importantly, all clones showed similar numbers of initial infections, as measured by the total number of infection sites and plaques (Figure 3C), indicating that viral entry was equally probable across clones. However, they differed dramatically in the subsequent spread of the virus. In the majority of clones (84.1%, n = 3), which we classified as “nonresistant,” the percentage of initial infections that went on to form plaques was around 85%, measured by computing the number of plaques divided by the number of plaques plus the number of non-progressing (nuclear only) infections. In contrast, a small subset of clones (15.9%, n = 3), which we classified as “resistant,” effectively contained the infection, meaning that the fluorescent reporter signal never left the nucleus (indicating no cell-to-cell spread) and only 2-20% of initially infected cells developed into plaques. This same phenomenon was observed in human primary keratinocyte clones, where 4.2% of clones demonstrated similar resistance to viral spread (n = 2) (Supplementary Figure 6A, B). The plaques that did form in the resistant clones were smaller than the ones forming in the nonresistant clones (Supplementary Figure 6C). Overall, these results are consistent with the lineage-encoded viral resistance that we observed in the organotypic skin constructs.

To determine the timescale of fluctuation driving this lineage-encoded viral resistance, we assessed resistance to viral spread in single-cell clones at various intervals after their initial isolation. We observed that after 3 weeks in culture, 2-20% of infection sites progressed in these resistant clones (Supplementary Figure 7A, B), but this number increased to around 80% after 5 weeks of culture, suggesting that these cells will eventually lose their resistance properties (Supplementary Figure 8A, B).

### Resistant clones have distinct chromatin accessibility profiles

It is often postulated that cellular memories are encoded in the structure or modification of chromatin. For the multi-omic characterization of resistant and nonresistant clones, we cultured single-cell clones, grew them to confluency, and split them into multiple parallel cultures for further characterization (Figure 3D). One split was infected with HSV-1 ICP4-YFP to categorize them as being resistant or nonresistant according to the criteria described earlier. The other splits were used for bulk ATAC sequencing and proteomics. This approach allowed us to compare the molecular profile of resistant vs. nonresistant clones in their initial, uninfected state.

Dimensionality reduction on ATAC-seq called peaks shows that the resistant clones separated from the nonresistant clones (Figure 3E), indicating broad differences in the different chromatin accessibility profiles of these two categories of clones. Nonresistant clones clustered closely to each other, while resistant clones appeared to be more heterogeneous. Hierarchical clustering of clones based on the pairwise Spearman correlation of the ATAC sequencing data, again calculated by binning reads, confirmed that result (Supplementary Figure 9A). Since clones R1 and R3 clustered closely in the PCA plot, we focused our downstream analysis on comparing R1 and R3 versus NR1–NR3, hypothesizing that R2 either started to lose its resistance phenotype, or its resistance is dictated by other epigenetic mechanisms. Differential peak analysis of R1 and R3 versus NR1–NR3 revealed 27,049 differentially accessible peaks in the resistant clones, compared to 18,209 in the nonresistant clones (Figure 3F).

### AP-1 binding sites are enriched in resistant clones

We then focused on particular motifs to try and identify transcription factors that may be responsible for memory encoding. We looked for motifs that were enriched in areas of differentially accessible chromatin across the resistant and nonresistant clones.

We quantified variations of all known motifs across samples using ChromVAR (for putative transcription factor activity) and performed de-novo motif discoveries using the differential peaks between resistant vs. nonresistant clones with HOMER (for motif identification within accessible peaks) [27]. Both analyses identified FOS:JUN motif accessibility as enriched in 2 (NR1 and NR3) out of 3 resistant clones, giving the highest overall deviation Z score of 43.407 (Figure 3G, Supplementary Figure 9B). Comparing the two FOS:JUN motif-enriched resistant clones to the nonresistant clones in our HOMER analysis revealed that AP-1 transcription binding motifs appeared in 61.56% of the differential peaks, compared to a 38.64% background (p < 0.001). To confirm whether this increased accessibility reflected actual transcription factor occupancy, we applied TOBIAS footprinting analysis [28]. With an overall differential binding score of 0.593 (p < 0.001), we infer that AP-1 binding sites are not only enriched in accessible chromatin regions in the resistant clones, but that AP-1 is also actively binding its target motifs.

To determine whether the predicted binding of AP-1 to target motifs is supported by changes in protein abundances, we performed quantitative mass spectrometry-based proteomic analysis of the same resistant and nonresistant clones. We focused our analysis on the resistant clones 1 and 2 (R1, R2), nonresistant clones 1, 2, and 3 (NR1–NR3), in parallel with analysis of the primary culture as control. Resistant clone 3 exhibited lower cell numbers and was hence retained for downstream analyses. Dimensionality reduction showed similar clustering of resistant and non-resistant clones as observed from the ATACseq data, confirming that alterations are also captured at the proteome level (Figure 3H). Numerous proteins were found to have differential abundances between resistant and non-resistant clones, with over 250 and 100 proteins having significantly higher expression in the resistant and nonresistant clones, respectively (Figure 3I). Among those most prominently regulated, we identified proteins associated with AP-1 signaling (e.g., APOE, CTHRC1), viral recognition (e.g., LOXL1, VIM), antiviral signaling (e.g., CD151, IFI44, ITGA6), and antiviral effector responses (e.g., SERPINB2, MMP1, TIMP1, TIMP3) (Supplementary Figure 10). Many of these proteins contain AP-1 binding sites, of which SERPINB2 is highlighted as an example (Figure 3J). AP-1 transcription factor JunB was detected at higher quantities in resistant clones versus nonresistant clones (adjusted p value of 0.0028), albeit at a subthreshold fold change of only 1.45.

Together, these multi-modal analyses identified AP-1 as a potential key transcriptional regulator associated with the resistant phenotype. These results are in line with recent work that has demonstrated an emerging role for AP-1 in epigenetic memory formation [29,30].

### Endogenous AP-1 upregulation induces viral resistance

The identification of AP-1 binding motifs as sites of potential regulatory differences between resistant and nonresistant clones led us to hypothesize that activation of endogenous AP-1 could make cells more resistant to viral infection. To test this hypothesis, we used Phorbol 12-myristate 13-acetate (PMA), a phorbol ester that activates protein kinase C, to activate AP-1 transcription factors [30]. We reasoned that, by treating cells with PMA before infecting them, an increased number of AP-1 transcription factors would bind and open chromatin near AP-1-associated genes, which in turn would make cells more resistant to viral replication.

We treated human primary skin fibroblasts and keratinocytes with PMA for various durations prior to infection with HSV-1 (Figure 4A). PMA pretreatment enhanced viral resistance drastically, both reducing the plaque percentage and plaque size (Figure 4B, Supplementary Figure 11A). PMA is known to activate nuclear factor-kappa B (NF-κB), thereby enhancing inflammatory responses, which could partially explain the observed reduction in plaque size—independent of AP-1’s role as a master regulator of cellular memory [31]. This is a possibility that we currently cannot rule out, and needs to be addressed by future research. Of note, the PMA-induced infection phenotype characterized by nuclear-restricted fluorescence is similar to that observed in resistant clones, supporting a causal role for AP-1 in mediating resistance memory.

**Figure 4:**
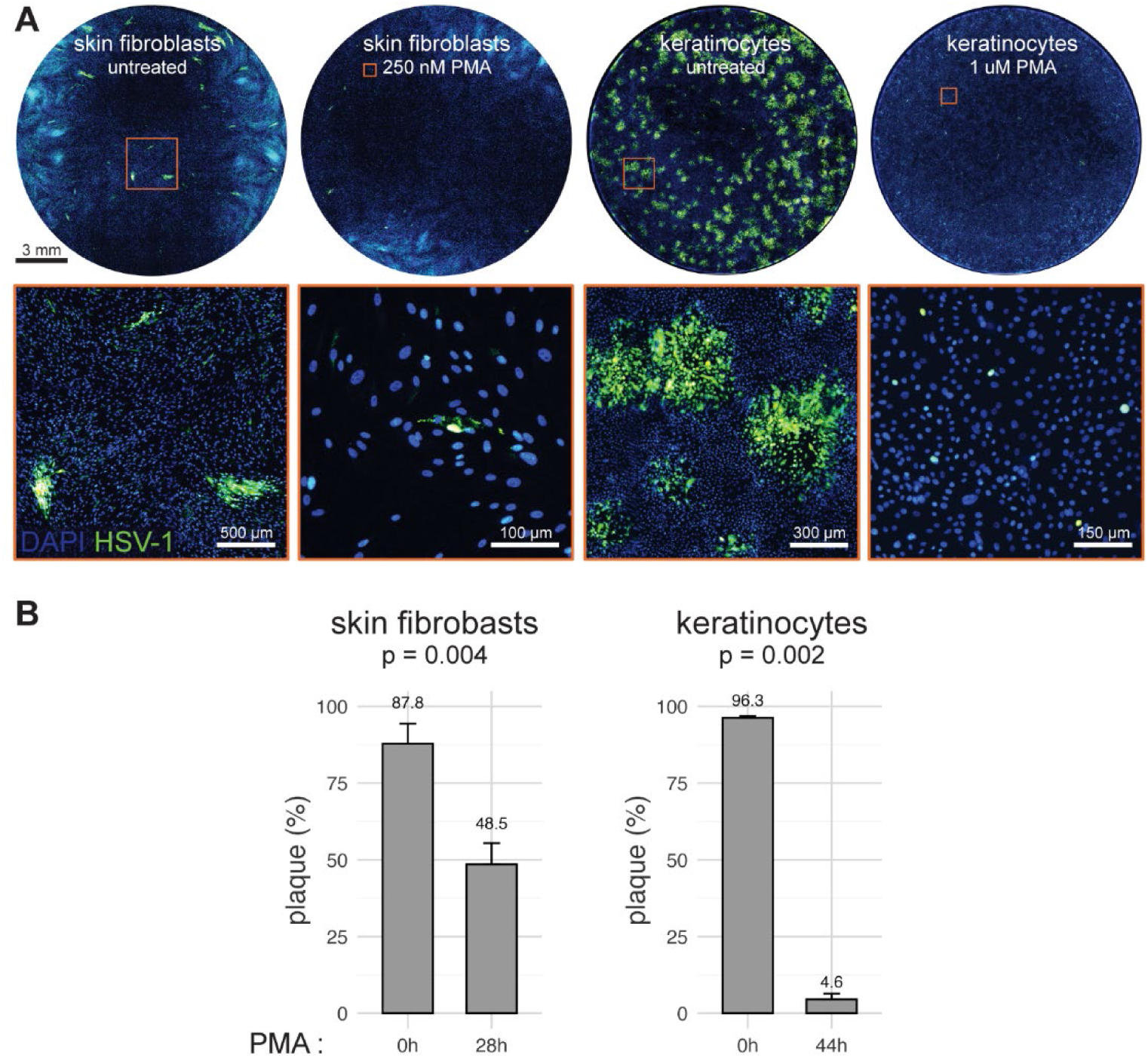
Endogenous upregulation of AP-1 induced resistance in human primary fibroblasts and keratinocytes. A) : Representative microscopy images of human primary skin fibroblasts and keratinocytes, untreated or treated with PMA, and infected with HSV-1 for 24 hours. Orange boxes indicate corresponding zoom in represented on the bottom row. B) : Bar graphs of mean + standard deviation and absolute mean of plaque percentage for untreated and PMA treated skin fibroblasts (n = 3) and keratinocytes (n = 2).

The timing of PMA administration turned out to be crucial, giving varying results for the two cell types tested (Supplementary Figure 11B). Administration of PMA 20 hours pre-infection was very effective in inducing the resistance phenotype in human primary keratinocytes, whereas for human primary fibroblasts, maximum effectiveness was reached at PMA administration of 4 hours pre-infection. In conclusion, these results suggest that AP-1 encodes lineage memory of viral resistance.

## Discussion

Clonal heterogeneity has emerged as a key driver of functional diversity in seemingly identical cell populations, shaping how cells respond to perturbations such as drugs, inflammatory signals, and viral infections. Seminal work by Shaffer et al. demonstrated that drug resistance in cancer can arise from rare pre-existing cell states, which can persist due to transcriptional variability and lineage-encoded memory, rather than stochastic noise [32]. Similarly, Clark et al. revealed clonal differences in NF-κB signaling, showing that only subsets of related cells mount strong transcriptional responses to inflammatory cues, reflecting clonal memory in pathway sensitivity [6]. Petkidis et al. further extended this principle to viral infections, where they found that clonal cell populations exhibit differential susceptibility to viruses [12]. They reported viral resistance in certain clonal lineages obtained from an immortalized cell line that was maintained for at least 9 weeks of cell cultivation. Here, we translate this concept to human primary cells in a tissue context, showing that lineage memory shapes viral resistance barriers in a human organotypic skin model. Together, these studies support the idea that functionally relevant lineage memory is not only limited to cancer biology, where it is studied most extensively, but is a generalizable principle observed across diverse biological systems, including antiviral immunity.

The functional consequences of different durations of lineage memory remain unknown. An attractive hypothesis is that memory can lead to very different spatial distributions in tissue. For instance, while IFN-I expression is limited to only a small fraction of infected or stimulated cells, the transient nature of the underlying memory ensures that IFN-I-producing cells are evenly distributed among tissues, with larger groups of related cells consistently including at least one IFN-I producer. This pattern aligns with the fate priming hypothesis, which proposes that subtle, sub-threshold fluctuations in key regulatory factors create a poised state in a subset of cells, enabling them to rapidly and robustly respond to stimuli when triggered [33]. In our system, we propose that a small, transiently primed subset of cells is responsible for initiating the IFN-I response upon infection, inducing ISG expression through autocrine and paracrine signaling. Since this priming state lasts only a few generations, it prevents clustering of high responders and instead favors a spatially distributed, fractional activation pattern.

In contrast to the fast-fluctuating IFN-I production memory, the viral resistance phenotype we describe appears to reflect a slow-fluctuating cell state that persists across many generations. This long-lived memory gives rise to large clusters of clonally related resistant cells within the tissue. Unlike the IFN-I response, which depends on rapid sensing and signaling, viral resistance in these clones appears to preexist prior to infection, suggesting a cell-intrinsic, epigenetically encoded state. Whether this state simply enables these cells to more rapidly initiate antiviral programs—and thereby more effectively limit viral replication and spread—remains to be fully elucidated. Differences in chromatin accessibility could underlie such shifts in response timing, consistent with principles of trained immunity. In the context of viral replication, responses are critical at the timescale of minutes to hours.

The observation that AP-1 motif accessibility and transcriptional activity was higher in resistant clones adds an additional mechanistic layer to this memory. The AP-1 family of transcription factors has previously been implicated in shaping cellular responses to inflammation, stress, and infection, often through epigenetically regulated enhancer programs. Recent work by Fuchs and colleagues demonstrated that AP-1 units JUN and FOS maintain residence on chromatin, thereby “indexing” regions in stress response genes that can be rapidly recruited during a secondary response for reactivation [29]. In a different context, Li et al. described how AP-1 mediates cellular adaptation and memory formation during therapy resistance [30]. In our context, AP-1 may function as a memory-inducing transcriptional regulator that helps imprint a resistant state across cellular generations, representing a form of innate immune memory encoded in chromatin accessibility.

Together, our findings highlight the importance of lineage in shaping immune responses, revealing that antiviral immunity is not solely governed by fast, dynamic fluctuations but also by long-lived clonal states. Understanding these distinct timescales of memory—transient priming for signaling and durable imprinting for resistance—may inform therapeutic strategies that harness or modulate cell-intrinsic immunity in both infectious and inflammatory disease contexts.

## Data and Code Availability

The raw and processed sequencing data generated in this study have been deposited in the NCBI Gene Expression Omnibus (GEO) under accession number GSE300510. The data can be accessed at https://www.ncbi.nlm.nih.gov/geo/query/acc.cgi?acc=GSE300510. The mass spectrometry proteomics data have been deposited to the ProteomeXchange Consortium via the PRIDE partner repository with the dataset identifier PXD065515. All raw and processed data as well as code for analyses performed in this study can be found using this link: https://www.dropbox.com/scl/fo/xseyvae1x6h8w8eq6akyq/AI0IZQkVtO6yMAboaQ0RMXs?rlkey=2k44xs0hv0pw7hsczajvyta4q&st=08yq9r9e&dl=0

## Acknowledgements

We thank the Raj lab members for scientific discussion and comments on the manuscript. In particular, we thank Pavithran Ravindran for their discussion on ATAC sequencing data analysis and Vinay Ayyappan for their support regarding the ATAC library preparation workflow. We thank Sydney Bracht and Staci Rakowiecki for their assistance with cryosectioning; the William Greenleaf lab for sharing their expertise in ATAC sequencing and providing us with the custom primers used in our ATAC-seq experiments; the Kavitha Sarma lab at The Wistar Institute for generously providing the Tn5 transposase; the Shaffer lab, in particular Sydney Shaffer and Christopher Cote for providing assistance and equipment for sequencing. Dermis for organotypic skin was provided by the Penn Skin Biology and Diseases Resource-based Center, funded by NIH/NIAMS grant P30-AR069589 and the University of Pennsylvania Perelman School of Medicine. This project has been made possible in part by grant 2023-332391 from the Chan Zuckerberg Initiative DAF, an advised fund of Silicon Valley Community Foundation. A.R. acknowledges support from a center grant from the Mark Foundation for Cancer Research, NIH Director’s Transformative Research Award R01 GM137425, NIH R01 CA238237, NIH R01 CA232256, and NIH 4DN U01 DK127405. A.S. acknowledges support from NIH-NIGMS through grant R35GM148351. I.C. acknowledges support from NIH NIAID AI174515 and NIH NIGMS GM114141.

## Author Contributions

L.E. and A.R. conceived and designed the project. L.E. designed and performed experiments with help from A.O and G.K., under supervision of A.R. G.K. optimized culturing organotypic skin and assisted in analysis and interpretation of the data. A.O. optimized the ATAC-seq protocol and assisted in analysis and interpretation of the data. J.Z. performed ATAC-seq analyses under supervision of K.Z. C.T. designed and optimized lineage labels. R.I. performed mass spectrometry and together with T.G. performed proteomic analysis under supervision of I.C. M.O. and N.D. provided virus stocks and helped interpreting the results. A.Sr. and A.Si. assisted in the sibling-cousin analysis and performed mathematical modeling for the k_on_ and k_off_ rates. L.E. and A.R. wrote the manuscript with input from all authors. All authors read and approved the final manuscript.

## Declaration of Generative AI and AI-Assisted Technologies

During the preparation of this work, the authors used ChatGPT and Claude to generate and improve code for analyses. The authors reviewed and edited the content and take full responsibility for the content of the manuscript.

## Materials and Methods

### Cell culture and viruses

Primary human keratinocytes were obtained from the Skin Translational Research Core (STaR) at the University of Pennsylvania, who isolated them from fresh surgical specimens. Cells were cultured in 50% Keratinocyte SFM medium (#17005042) and 50% 154 medium (#M154500), supplemented with 750 pg/mL recombinant epidermal growth factor (EGF), 15 ug/mL bovine pituitary extract (BPE), and 5% penicillin/streptomycin (Invitrogen, #15140122). Cells were passaged every 2-3 days using 0.05% trypsin-EDTA (Invitrogen, #25300054). Primary human skin fibroblasts were obtained from commercial sources, obtained from neonatal male donors (Cascade Biologics, #C0045C; ATCC, #CRL-2522). Cells were cultured in DMEM (ThermoFisher, #11995065), supplemented with 10% fetal bovine serum (FBS) and 5% penicillin/streptomycin (Invitrogen, #15140122). Cells were passaged every 3-4 days using 0.05% trypsin-EDTA (Invitrogen, #25300054). Cell cultures tested negative for mycoplasma.

Recombinant vesicular stomatitis virus VSV carrying the M51R point mutation, generated by Dr. Sean Whelan [34,35], were a kind gift from Megan H. Orzalli at the University of Massachusetts Chan Medical School. HSV-1 (strain 17) viruses expressing ICP4-YFP, generated by Roger Everett [36], were a kind gift from Nir Drayman at the University of California, Irvine. Viral stocks were prepared, propagated and titrated in Vero cells according to well-established protocols [37]. *In vitro* infections were performed by incubating small volumes of virus inoculum on cell monolayers for 1 hour, after which all inoculum with non-infected virus particles got removed. Next, cells were cultured in a high viscous medium containing 1.5% methylcellulose to avoid secondary infections through diffusion.

### Organotypic skin cultures

Organotypic skin cultures were made using well-established protocols [16,17]. In short, devitalized human acellular dermis, obtained from the University of Pennsylvania’s Skin Biology and Diseases Resource-based Center, was used as a supporting extracellular matrix on top of a 3D printed stand to allow human primary keratinocytes to be cultured in an air-liquid interface. Keratinocytes were seeded onto the basement membrane side.

### Lineage labeling

To generate the nuclear-localized fluorescent reporter constructs, we cloned eGFP, mOrange, and mKate2 fluorescent proteins into a lentiviral backbone under the control of the EF1α promoter, similar to what has been described previously [38]. Fluorescent proteins were flanked on both sides by the SV40 nuclear localization signal (NLS). The constructs included the woodchuck hepatitis virus posttranscriptional regulatory element (WPRE) to enhance transcript stability and transgene expression. A β-lactamase (AmpR) resistance gene was used for bacterial selection during plasmid amplification.

Plasmid amplification was performed upon plasmid electroporation into Endura electrocompetent Escherichia coli cells (Lucigen) using a Gene Pulser Xcell (Bio-Rad). A AmpR resistant clone was isolated and cultured on a shaker at 32 C for 12-14 hours. Cells were pelleted, and plasmids were isolated using the EndoFree Plasmid Maxi Kit (Qiagen, 12362) according to the manufacturer’s protocol.

Lentivirus was produced by transfecting Lenti-X 293T cells with helper plasmids VSVG and psPAX2 using polyethylenimine (Polysciences 23966) in Opti-MEM (Thermo Fisher, #31985062). A mass ratio of 4:2:3 for plasmid DNA:VSVG:psPAX2 was used. Transfection mix was gently mixed, incubated for 15 minutes at room temperature, before adding dropwise to the cell monolayer. Lentivirus was collected 24, 48 and 72 hours after transfection, pooled, sterile filtered, and concentrated using ultracentrifugation. Lentiviral stocks were stored at −80 C in PBS containing 2.5% glycerol.

### RNA FISH

We designed custom oligonucleotide probe sets complementary to our genes of interest using a custom MATLAB software (available at https://flintbox.com/public/project/50547/). Oligonucleotides were ordered from IDT (probe sequences available in Supplemental Table 1). We pooled 15-32 oligonucleotides per gene, after which an amine group was added on the 3’ end with TdT and coupled to a fluorophore (Cy3, Alexa 594, Atto647N, or Atto700). Single-molecule RNA FISH was performed as earlier described [39]. In short, we fixed cells and tissue sections in 4% formaldehyde solution for 10 minutes and permeabilized them with 70% ethanol for an hour, both at room temperature. We next washed the cells and tissue sections with washing buffer containing 10% formamide and 2x SSC. For hybridization, we applied hybridization buffer containing the RHA FISH probes and 10% formamide, 2x SSC, and 10% dextran sulfate, incubating overnight at 37 C. Before imaging, the samples were washed twice with washing buffer. For imaging, samples were stained with DAPI and transferred to 2x SSC.

Samples were imaged on a Nikon Ti-E with a 60X Plan-Apo objective and filter sets for DAPI, Cy3, Atto647N, Alexa594, and Atto700. Nikon Elements imaging software was used to stitch multi-tiled images. The Nikon Perfect Focus System ensured images remained in focus. All images used for gene transcription quantification were acquired at 60x magnification and 1.4 numerical aperture.

### Imaging and analyses

We analyzed our image datasets using the NimbusImage platform, which is available here: https://github.com/arjunrajlaboratory/NimbusImage/ and hosted here: https://www.nimbusimage.com/. We used the Cellpose and Cellpose retrain tools [40–42] to detect cells and the Piscis and Piscis retrain tools [43] to detect mRNA spots. We connected each mRNA spot to the nearest cell using the “Connect to nearest tool” as described in the NimbusImage documentation.

### Viral infections in the organotypic skin

To introduce viral infections in the organotypic skin, we opted to use a well-established, reproducible method [14]. In short, full-grown tissue was removed from the organotypic stand and placed epidermis facing up on a flat sterile surface. A commercially available microneedling device (roller with needles of 1.5 mm in length and 0.25 mm in diameter, Amazon) was used to perforate an even distribution of micro holes by rolling it over the tissue back and forth multiple times, in multiple directions. The perforated tissue was placed back on the organotypic stand before pipetting a small volume of highly concentrated virus inoculum (∼1×10^6 PFU in 20 μL) on top of the epidermis. After 3 days of incubation, tissue was fixed and embedded in Tissue-Tek® O.C.T. Compound (Sakuraus, #4583) for cryosectioning.

### Clone analysis in the organotypic skin

In organotypic skin, clones were manually annotated as horizontal blocks oriented parallel to the basement membrane, based on the expression of the nuclear fluorescent lineage labels. Clones were annotated if they included at least 3 cells of the same color, with only a maximal distance of 80 µm between individual cells. Additional manual annotations included regions of VSV replication, and the total imaged epithelial region, all aligned in the same orientation relative to the basement membrane. To assess spatial relationships, the median distance between each VSV-infected region and the nearest clonal boundary was calculated along the x-axis. The full annotated region was used to define the physical horizontal boundaries (x-coordinates) within which infected regions were computationally shuffled, while keeping the clone annotations fixed in their original positions. Shuffling was performed by randomly reassigning the horizontal location of the largest infected region first, followed by the second largest, and so on, ensuring no overlaps and maintaining original region sizes. This procedure was iterated 500 times to generate a distribution of permuted datasets for statistical comparison to the observed experimental data.

### Sibling and cousin analysis and modeling

In viral infection experiments *in vitro*, siblings and cousins were manually annotated, based on the expression of the nuclear fluorescent lineage labels. Siblings were annotated when exactly two cells of the same lineage were observed in close proximity, defined as within 500 μm of each other. Cousins annotated when four or more cells of the same lineage were present in spatial proximity. In cases of more than four cells present, the four nearest cells were selected for annotation as cousins, under the assumption that spatial proximity reflects more recent common ancestry following cell division. All cousin data represents all 6 possible combinations of the 4 individual cells. Neighboring cells were annotated independently of lineage, by manually selecting cells in spatial sequence based solely on proximity, without considering fluorescent labels.

For both viral susceptibility and IFN-I production—which were modeled as independent processes—we used a two-state stochastic switching model in which cells can transition between an ON state (susceptible or IFN-I producing, respectively) and an OFF state (resistant or non–IFN-I producing, respectively). Transitions occur from the ON state to the OFF state at a rate *k*_*OFF*_, and from the OFF state to the ON state at a rate *k*_*ON*_. In both states, cells proliferate with rate *λ*.

The following assumptions are made for this model:

1. The time a cell stays in the ON state is an independently and identically distributed (iid) random variable following an exponential distribution with mean 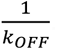.
2. Similarly, the time spent in the OFF state is an iid random variable following an exponential distribution with mean 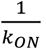.
3. The cell-cycle time is also an iid random variable following an exponential distribution with mean 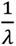.
4. Cell states are preserved during division, i.e., both daughters just after division inherit the same state of the mother. With this approximation, state switching only occurs during the interphase period.

At steady state, the average fraction of cells in the ON state are:

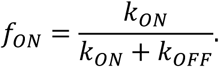

The probability that two daughters are both in the ON State is:

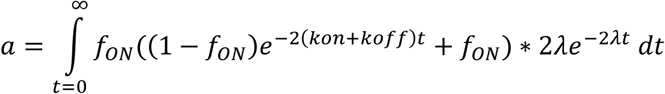

and evaluating the integral gives:

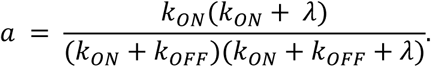

Similarly, the probability that two daughters are in the OFF State is:

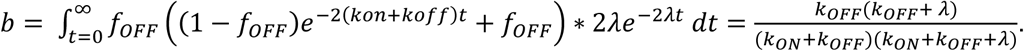

Finally, the probability of having daughters in different states is:

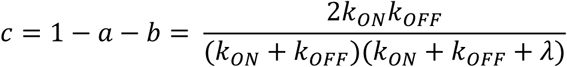

that yields the Odds Ratio defined as:

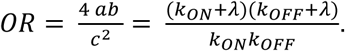

As expected, in the limit of fast switching that corresponds *k*_*O*_*_N_*, *k_*O*FF_* ≫ *λ, OR* converges to one, which corresponds to a purely random assignment of cell states.

For each set of pairs (siblings, cousins, and neighbors), two statistics are computed: the Odds Ratio and the probability of cells being in the ON state. An Odds Ratio of 1 indicates no memory (i.e., the fate of one cell does not influence the other), while a value greater than 1 indicates memory (i.e., the fate of one cell influences the other). A higher Odds Ratio indicates stronger memory between cells. The probability of cells being in the ON state is approximated by the ratio of the number of cells in the ON state (*N_*O*N_*) to the total number of cells (*N*):

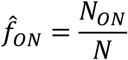

The Odds Ratio is calculated through the probabilities of joint states in a pair of cells. These are approximated using the fraction of pairs of cells in ON state (*â*), fraction of pairs of cells in OFF state (*b̂*), or fraction of pairs with one cell in ON state and one cell in OFF state (*ĉ*).

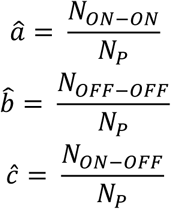

where *N_*O*N_*_−*ON*_ is the number of pairs with both cells in the ON state, *N*_*OFF−OFF*_ is the number of pairs with both cells in the OFF state, and *N*_*ON−OFF*_ is the number of pairs with one cell in the ON state and one in the OFF state. *N*_*P*_ is the total number of cell pairs. Then, the Odds Ratio is calculated experimentally as:

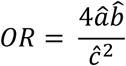

These values are then used to estimate the switching rates between the ON and OFF states by solving the equations:

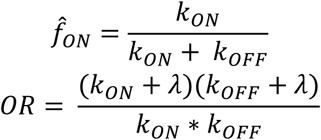

where *λ* is the cell proliferation rate, assumed to be 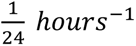. Bootstrapping was performed in two stages to estimate key parameters. In the first stage, resampling was conducted 1,000 times to calculate the probability of cells being in the ON state and the Odds Ratio for each resample. From these 1,000 bootstrapped samples, the mean *f_*O*N_* and mean OR were computed, along with 95% confidence intervals. In the second stage, these bootstrapped estimates were used to calculate the switching rates *k*_*ON*_ and *k*_*OFF*_, again, with1,000 bootstrapped resamples. The mean switching rates and their corresponding 95% confidence intervals were then obtained from this second round of bootstrapped samples. All analyses were conducted using R and RStudio (version 4.3.2)

Finally, the memory half-life—defined as the number of cellular divisions (roughly equal to days) required for a 50% probability of fate switching—was calculated using the following equations:

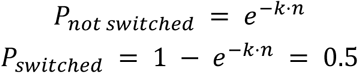

Setting *P_switched_* = 0.5 gives:

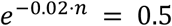

Taking the natural logarithm of both sides yields:

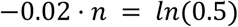

Solving for *n* gives the memory half-life:

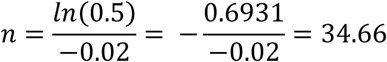

### ATAC sequencing analysis

We performed ATAC sequencing according to well-established protocols [44]. In short, we lysed cells and set up the transposition reaction with the Tn5 transposase at 37 C for 30 minutes. Next, we isolated transposed and adaptor-ligated genomic DNA with the Zymo DNA Clean and Concentrator Kit (Zymo, #D4014). We amplified the library using custom-made primers (Supplemental Table 2), originally designed by the William Greenleaf lab at Stanford University. The library was sequenced using AVITI Cloudbreak FS High Output 2×75 sequencing kit (#60-00015, Element Biosciences), on a AVITI sequencer (Element, Biosciences). 838.28 million reads were acquired and aligned to the hg38 assembly using bowtie2. Duplicates were removed, reads were shifted and normalized based on counts per million. Peaks were called using MACS2. Motif analyses (ChromVAR and HOMER) were performed on differential peaks. TOBIAS footprinting was performed on merged data of the 4 different groups: 1) both duplicates of the primary culture, 2) all 3 nonresistant clones and their duplicates, 3) 2 resistant clones and their duplicates, 4) 1 resistant clone and its duplicates.

### Proteomics analysis

Cells were harvested, pelleted, and washed twice with 1X PBS. A total of 100,000 cells per sample were lysed in 10% SDS (Sigma-Aldrich, #05030), and lysates were subsequently diluted to a final concentration of 5% SDS in 50 mM ammonium bicarbonate (Sigma-Aldrich, #A6141). Lysed samples were briefly sonicated prior to reduction and alkylation at 95°C for 5 minutes. Next, the soluble protein fraction was recovered and sample clean-up and protein digestion with trypsin was performed using S-Trap micro spin columns, according to the manufacturer’s instructions (Protifi, #C02-micro-80). Following digestion, peptides were sequentially eluted in 40 μL of (1) 25 mM TEAB, (2) 0.2% formic acid, and (3) 50% acetonitrile with 0.2% formic acid. Peptide eluates were pooled, dried using a vacuum concentrator, and resuspended in 4% acetonitrile containing 0.1% formic acid and 0.015% n-Dodecyl β-D-maltoside (DDM) prior to LC-MS/MS analysis.

Peptide samples were analyzed using a timsTOF Ultra mass spectrometer (Bruker) equipped with a CaptiveSpray 2 ion source and a 10 μm emitter (Bruker). Chromatographic separation was performed on a nanoElute 2 nanoLC system (Bruker) by direct injection of samples (150 ng) onto a PepSep C18 analytical column (1.5 μm particle size, 75 μm × 25 cm; Bruker). Peptides were eluted over a 60-minute linear gradient from 3% to 34% solvent B (0.1% formic acid in 99.9% acetonitrile), with solvent A consisting of 0.1% formic acid in water, at a constant flow rate of 200 nL/min. Intact peptide and fragment ion measurements were acquired using a data independent acquisition-parallel accumulation serial fragmentation (diaPASEF) method [45]. diaPASEF isolation windows were optimized using pydiaid [46], followed by manual adjustment of ion mobility boundaries.

Raw data from DIA acquisitions were analyzed using the FragPipe pipeline (v23.0, https://fragpipe.nesvilab.org/) with the default “DIA_SpecLib_Quant_diapasef” workflow [47]. Briefly, the workflow extracted pseudo-MS/MS spectra using diaTracer [48], which were directly searched by the MSFragger algorithm against the UniProt-SwissProt human and HSV-1 sequence databases (2025-03) supplemented with common contaminants. The peptide sequence matches were re-scored by MSBooster [49] using deep learning-based prediction models of MS/MS fragmentation (Prosit_2023_intensity_timsTOF), retention time (DIA-NN), and ion mobility (DIA-NN). The resulting peptide spectrum matches were analyzed by Percolator [50] and ProteinProphet/Philosopher [51,52] to control FDR to 1% at the ion and protein levels, respectively, using a target-decoy approach. The FDR-controlled precursor ions were assembled into a DIA spectral library by EasyPQP (v1.50.2). The DIA library was passed to DIA-NN (v2.1.0) [48] to perform quantitative analysis using the default FragPipe settings, except with the following additional parameters were specified as command line options: -- window 11 --mass-acc 15 --mass-acc-ms1 15.

The resulting “report.matrix_pg.tsv” file was processed using R to perform the following main steps: (1) assigned samples to resistant, non-resistant, and control groups, (2) filtered to retain protein groups with at least 25% valid values across all samples and at least 75% valid values in at least one sample group, (3) defined a function to perform imputation within each sample group when valid values were less than 2, using a random value selected from a normal distribution centered at the lower 1% abundance threshold, and (4) performed pairwise differential analysis using limma (ver. 3.62.2). The Benjamini-Hochberg (BH) procedure was used for multiple hypothesis test correction. Protein groups with an adjusted p-value ≤ 0.05 and absolute log₂ fold change ≥ 1 were considered differentially regulated.

### Statistical analysis

For each experiment, the sample sizes and number of replicates are indicated in the main text and figure legends. Statistical tests for calculating p values are indicated in the figure legends. All statistical analyses were performed in R unless otherwise stated.

## Supplemental Information

**Supplementary Figure 1:**
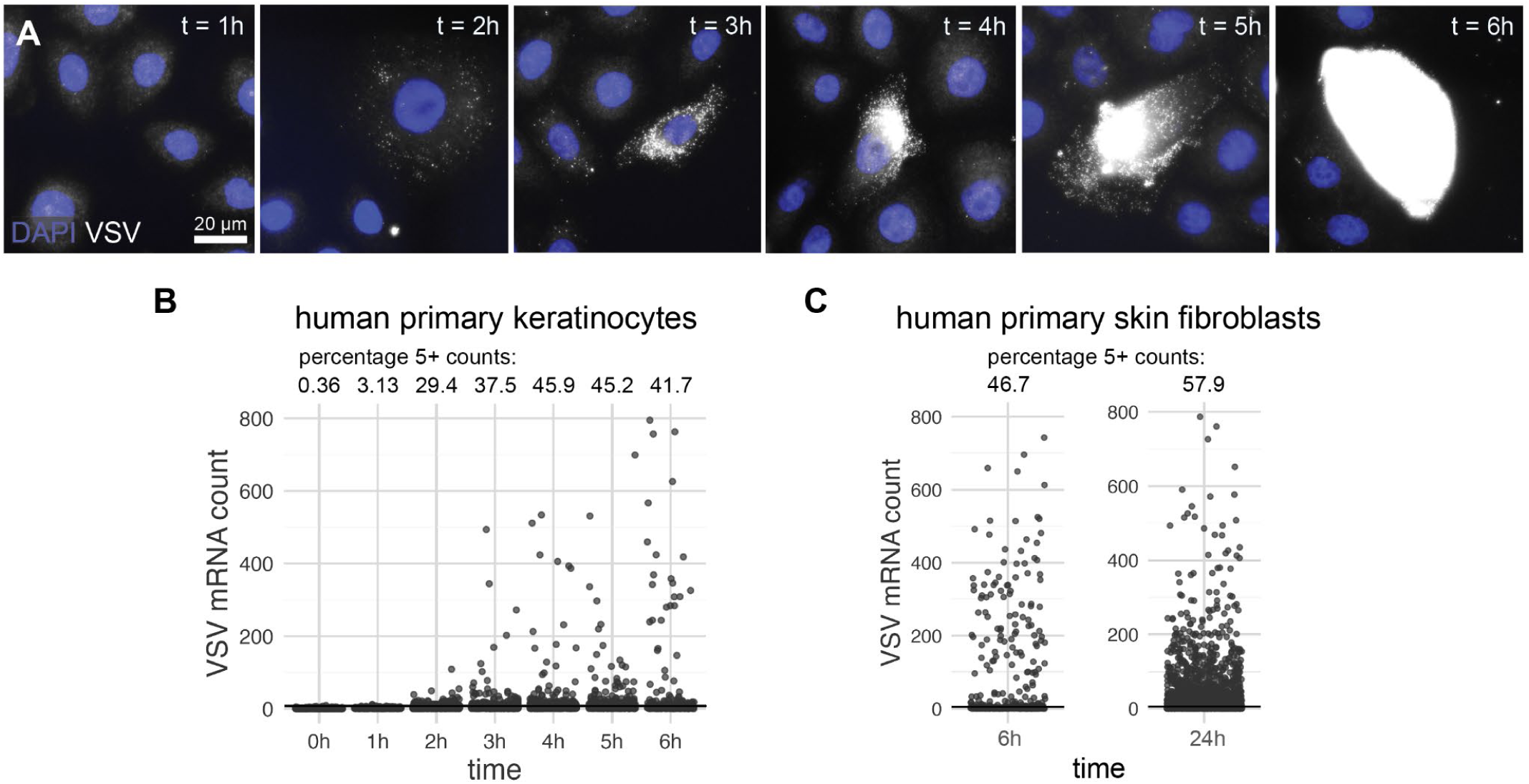
VSV replication happens fast, infecting roughly half of the primary fibroblasts after 24 hours. A) : Representative microscopy images of timelapse of human primary keratinocytes were infected with VSV at MOI = 2. B) : Quantification of VSV mRNA counts of timelapse presented in panel A. Black line indicates the threshold used for calculating the percentage of infected cells, n = 2. C) : Quantification of VSV mRNA counts in human primary fibroblasts infected with VSV at MOI = 2 for 6 and 24 hours. Black lines indicate the threshold used for calculating the percentage of infected cells, n = 2.

**Supplementary Figure 2:**
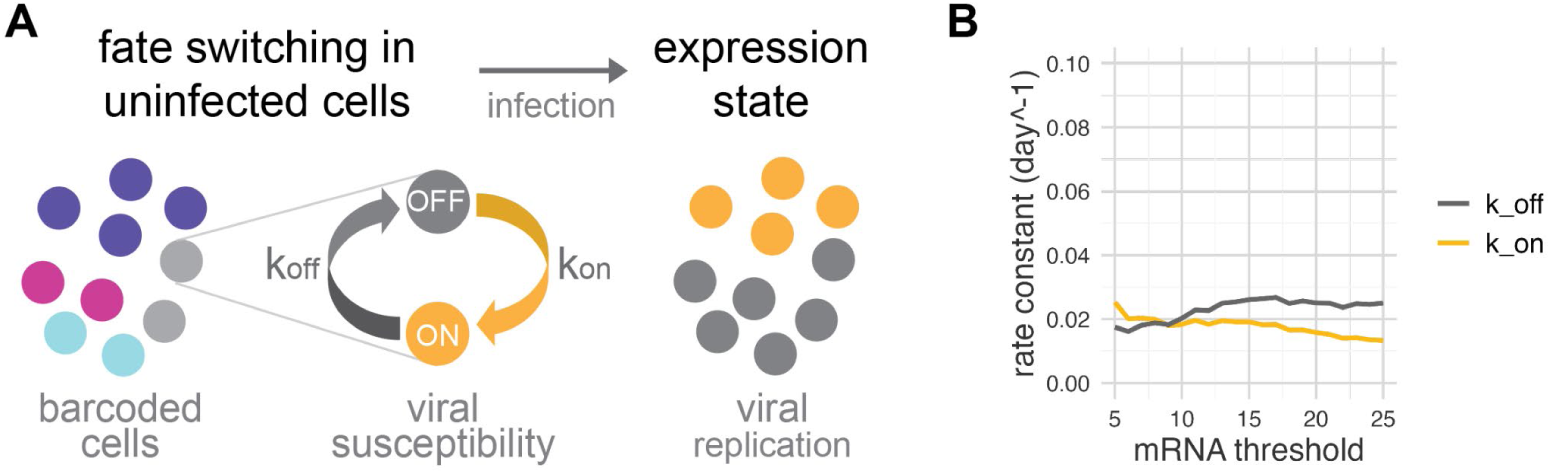
Viral susceptibility fate switching modeled with k_on_ and k_off_ rates. A) : Schematic on fate switching model. Before infection, cells switch between viral susceptibility states (ON and OFF), at rates k_on_ and k_off_. Cell lineages are tracked using barcoding approaches such as lineage labeling. Upon infection, these states are reflected by the viral RNA expression states, correlating with viral replication rates. B) : k_on_ and k_off_ rates at varying VSV mRNA thresholds based on experimental data presented in Figure 2.

**Supplementary Figure 3:**
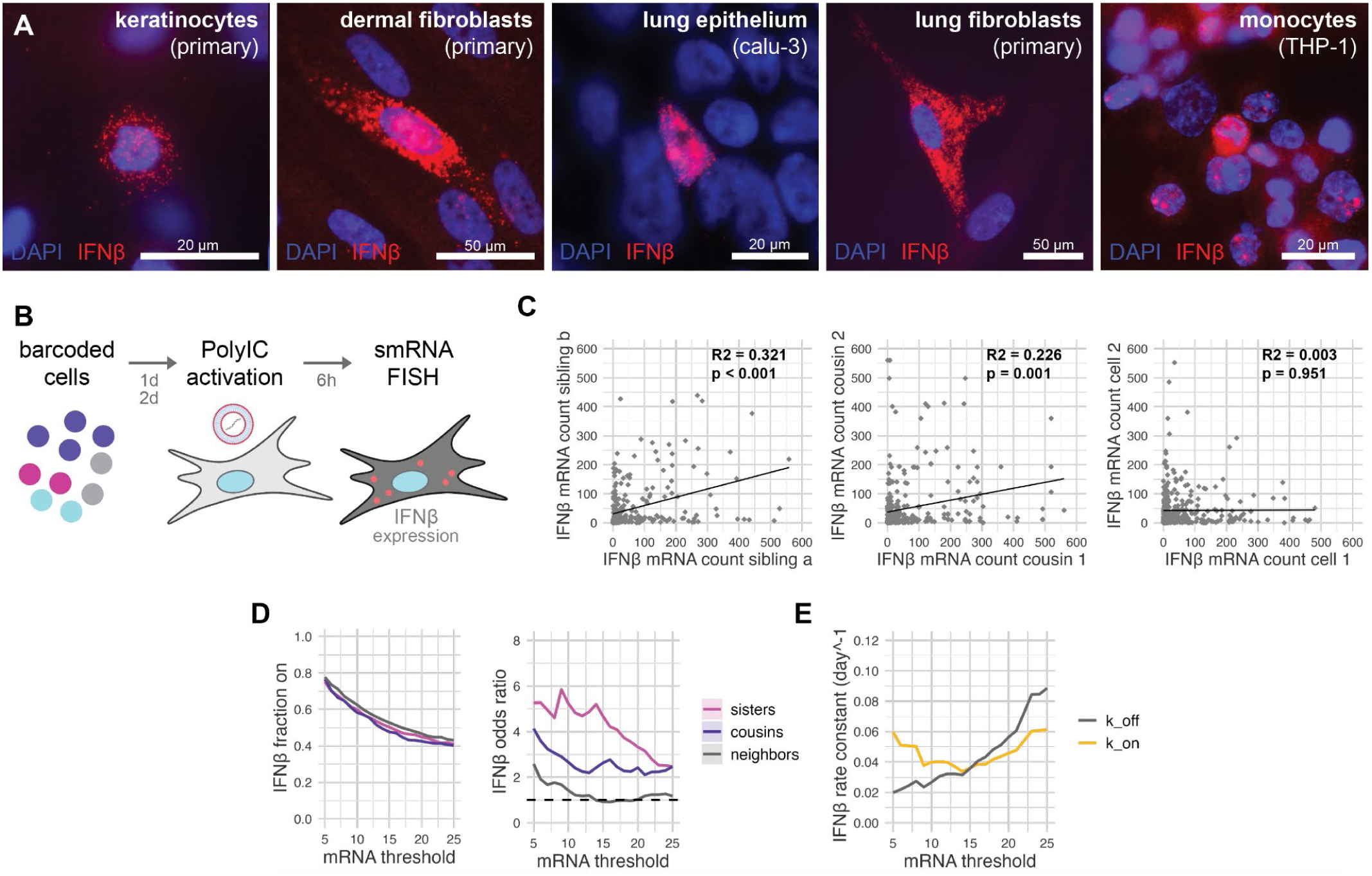
IFNβ memory is fast-fluctuating–PolyIC activation. A) : IFNβ production is highly heterogeneous, observed in human primary keratinocytes, skin fibroblasts, lung epithelium, lung fibroblasts and monocytes. Cells were activated with PolyIC for 6h. B) : Schematic representation of experimental setup. C) : IFNβ correlation plots for human primary keratinocyte siblings, cousins and neighbors (n = 3). D) : Line plots representing the different fractions on and the odds ratios for various mRNA thresholds. E) : k_on_ and k_off_ rates at varying IFNβ mRNA thresholds.

**Supplementary Figure 4:**
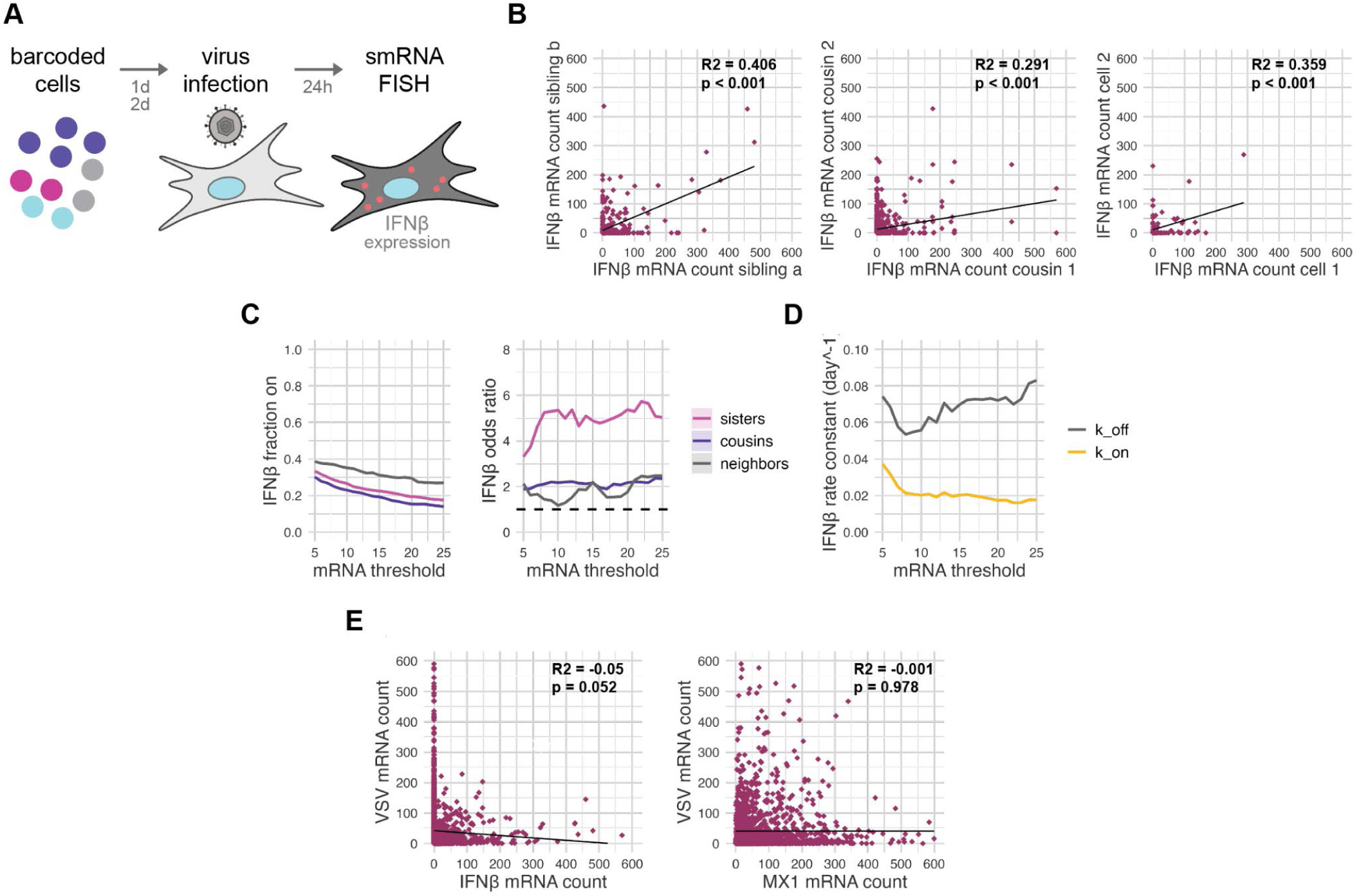
IFNβ memory is fast-fluctuating–VSV infection. A) : Schematic representation of experimental setup. Cells were infected with VSV for 24 hours. B) : IFNβ correlation plots for human primary skin fibroblast siblings, cousins and neighbors (n = 2). C) : Line plots representing the different fractions on and the odds ratios for various mRNA thresholds. D) : k_on_ and k_off_ rates at varying IFNβ mRNA thresholds. E) : Correlation plots comparing VSV mRNA counts with IFNβ and MX1 (n = 2).

**Supplementary Figure 5:**
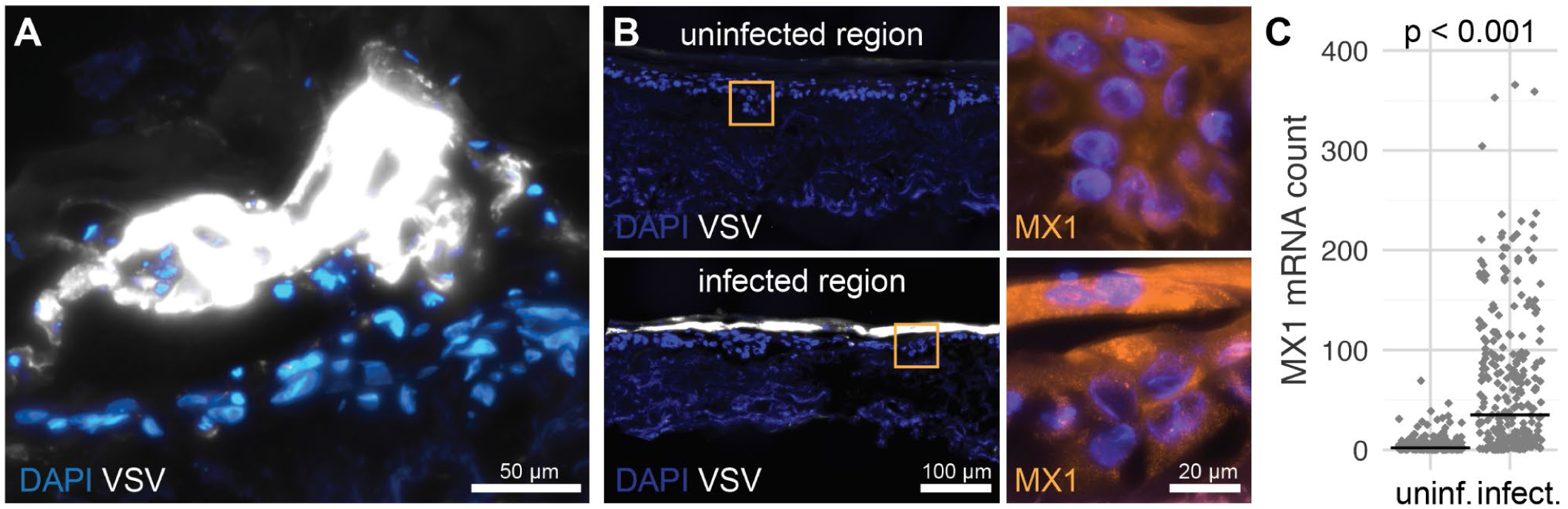
VSV infection in organotypic skin creates distinct replication zones with sharp boundaries and induces MX1 expression in adjacent uninfected cells. A) : 60x magnification of VSV-infected region. Organotypic skin was infected with VSV-M51R for 3 days. Viral replication was quantified with smRNA FISH. B) : Representative microscopy images of uninfected regions and infected regions, obtained from the same organotypic skin tissue. Orange boxes highlight zoom-in images with MX1 smRNA FISH signal. C) : Quantification of MX1 mRNA counts in uninfected regions versus infected regions. Welch Two Sample t-test, n = 3 biological replicates.

**Supplementary figure 6:**
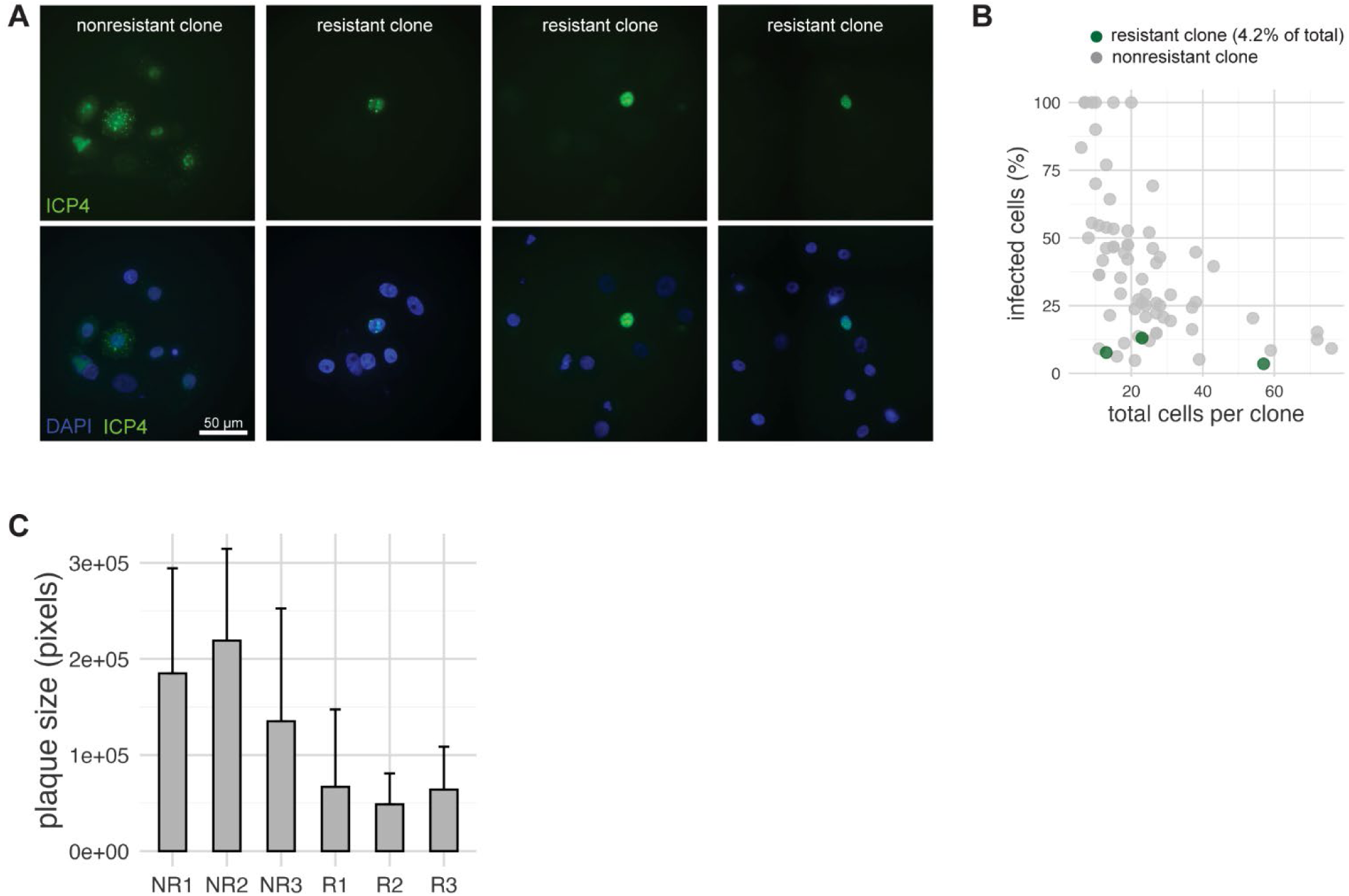
Human primary keratinocyte clones challenged with HSV-1 WT and plaque sizes in fibroblast clones. A) : Representative examples of nonresistant and resistant clones, infected with HSV-1 ICP4-YFP for 24 hours. Clones were annotated based on cell clusters generated upon low cell seeding. Clones were infected at rates of ∼2 virus particles per clone, calculated based on clone area. B) : Quantification of percentage of infected cells per clone, grouped by resistant and nonresistant clones. C) : Bar graph of mean + standard deviation of plaque size per nonresistant and resistant fibroblast clones.

**Supplementary Figure 7:**
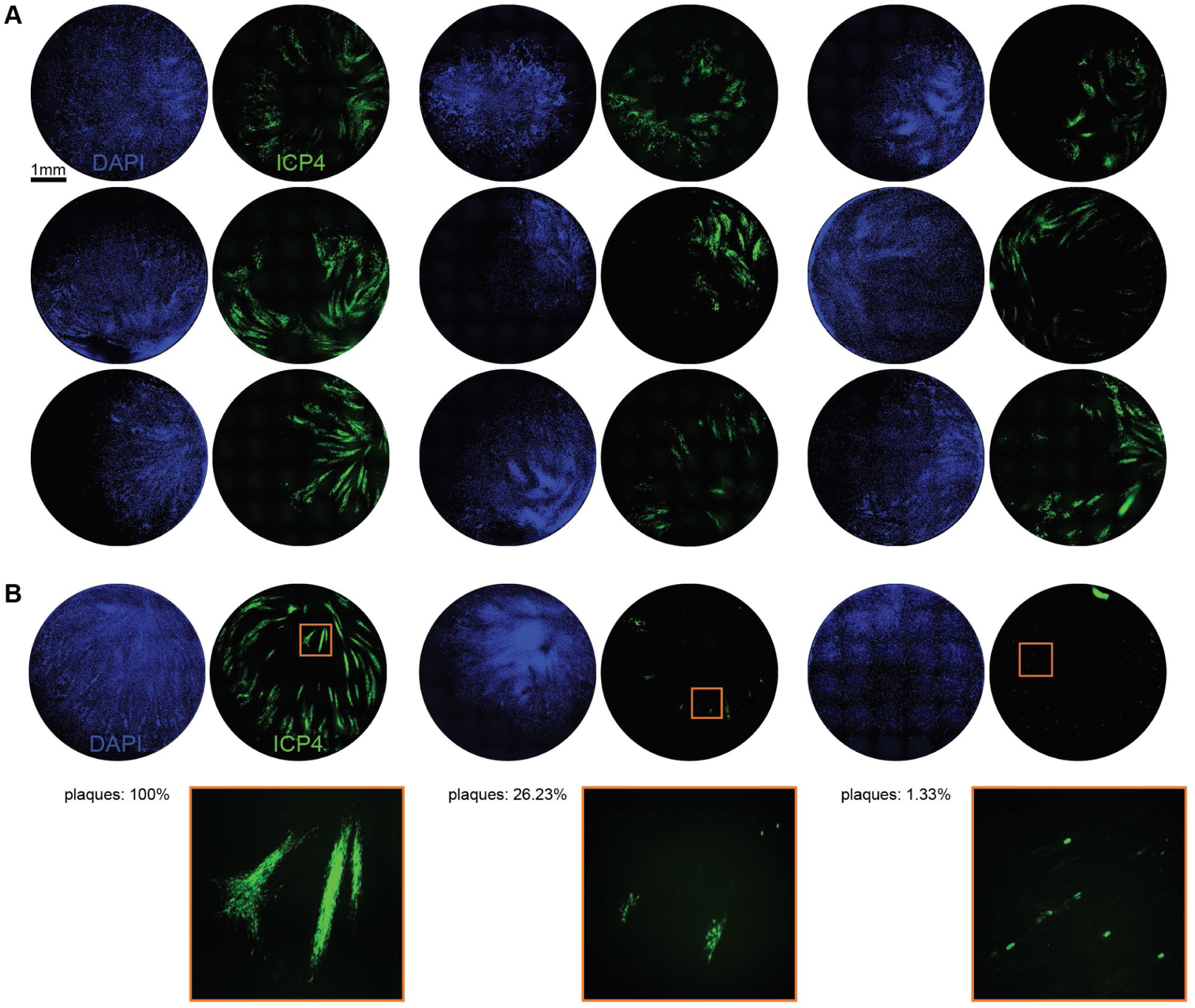
Representative examples of human primary fibroblast clones challenged with HSV-1 WT early after single-cell seeding. A) : Representative examples of nonresistant primary fibroblast clones challenged with HSV-1 WT 3 weeks after single-cell seeding. B) : Additional representative examples of a nonresistant clone and two resistant clones, as in panel A. Orange boxes indicate the corresponding zoom in. Indicated are the percentage of plaques over the total amount of infection sites and plaques.

**Supplementary Figure 8:**
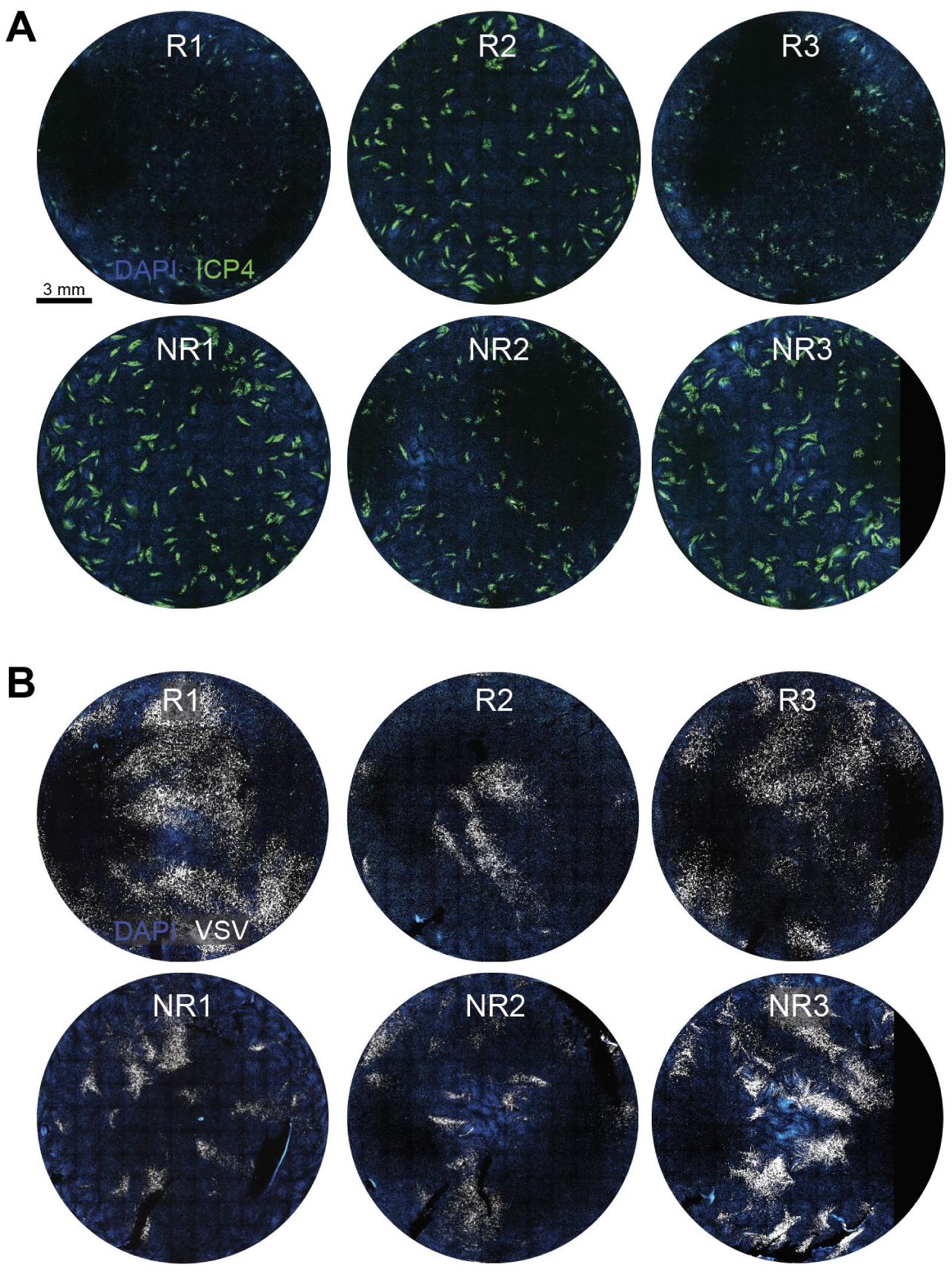
Representative examples of human primary fibroblast clones challenged with HSV-1 WT and rVSV-M51R, 8 weeks after single-cell seeding. A) : Resistant and nonresistant clones, corresponding to the clones characterized with ATACseq and proteomics, challenged with HSV-1 WT 8 weeks after single-cell seeding. B) : Resistant and nonresistant clones, corresponding to the clones characterized with ATACseq and proteomics, challenged with rVSV-M51R, 8 weeks after single-cell seeding.

**Supplementary Figure 9:**
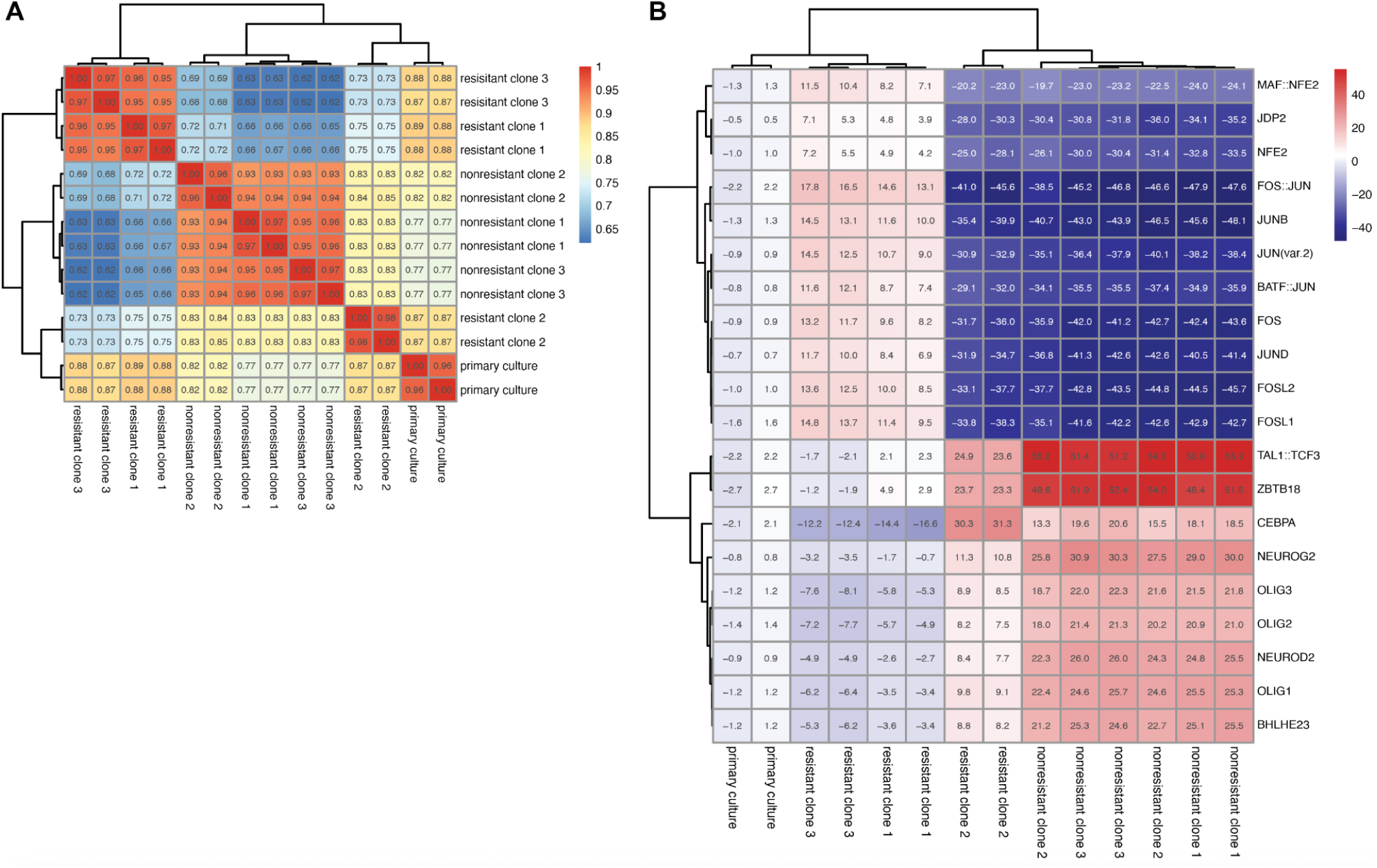
ATACseq–based clonal clustering and motif deviation analysis in resistant and nonresistant clones. A) : Hierarchical clustering of clones based on the pairwise Spearman correlation of the ATAC sequencing data, as calculated by binning reads. B) : Top 20 transcription factor motifs ranked by ChromVAR deviation Z-scores across resistant and nonresistant clones. Each row represents a motif, and columns represent individual clones. Higher Z-scores indicate increased motif accessibility.

**Supplementary Figure 10:**
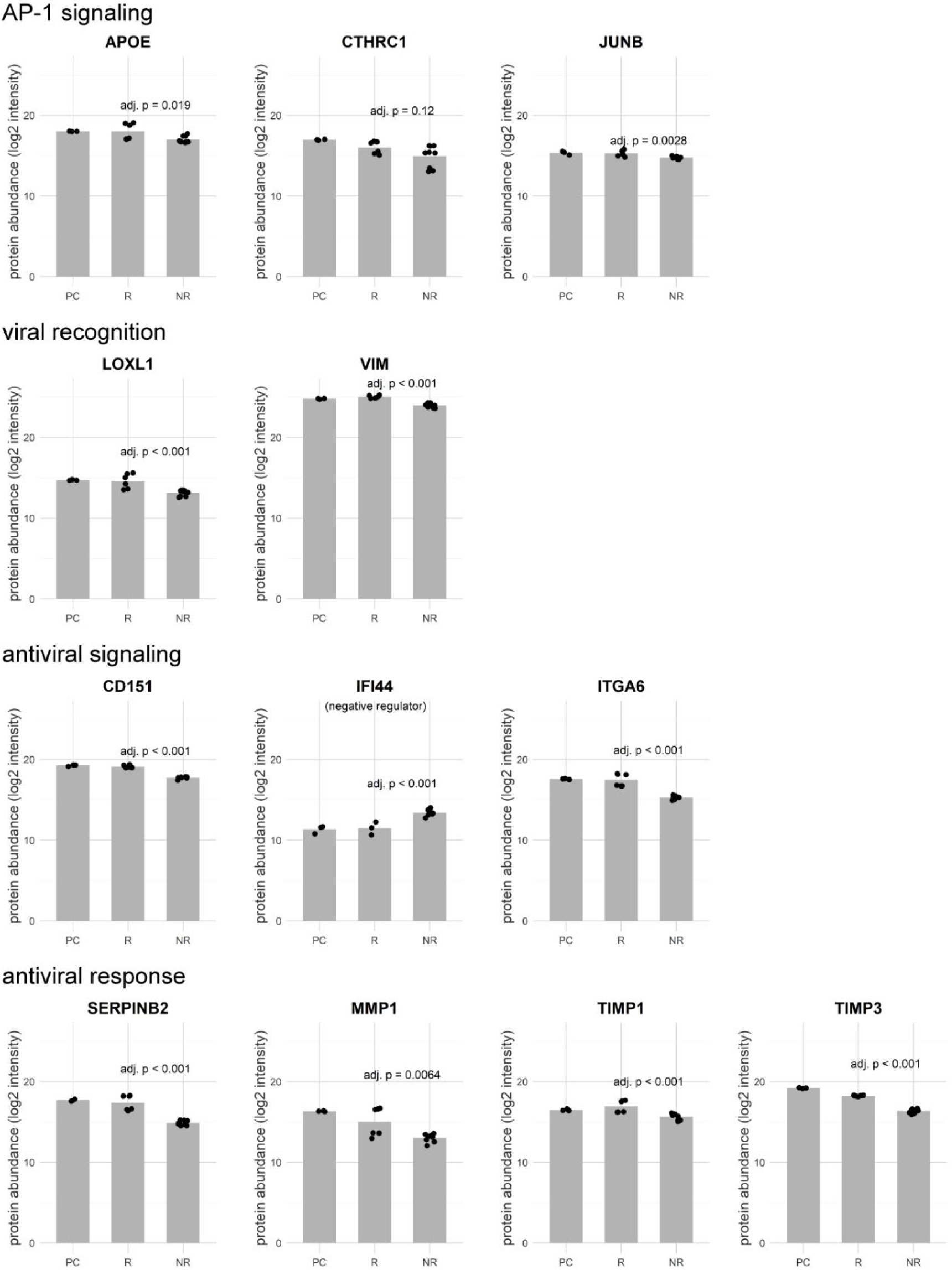
Proteomics data supports decreased AP-1 signaling and increased viral recognition, antiviral signaling and antiviral responses in resistant clones compared to nonresistant clones. Proteomics data on proteins related to AP-1 signaling, viral recognition, antiviral signaling, and antiviral responses. Adjusted p-values reflect statistical differences between R and NR conditions. Bars represent the means. Abbreviations: PC = primary culture, R = resistant, NR = nonresistant.

**Supplementary Figure 11:**
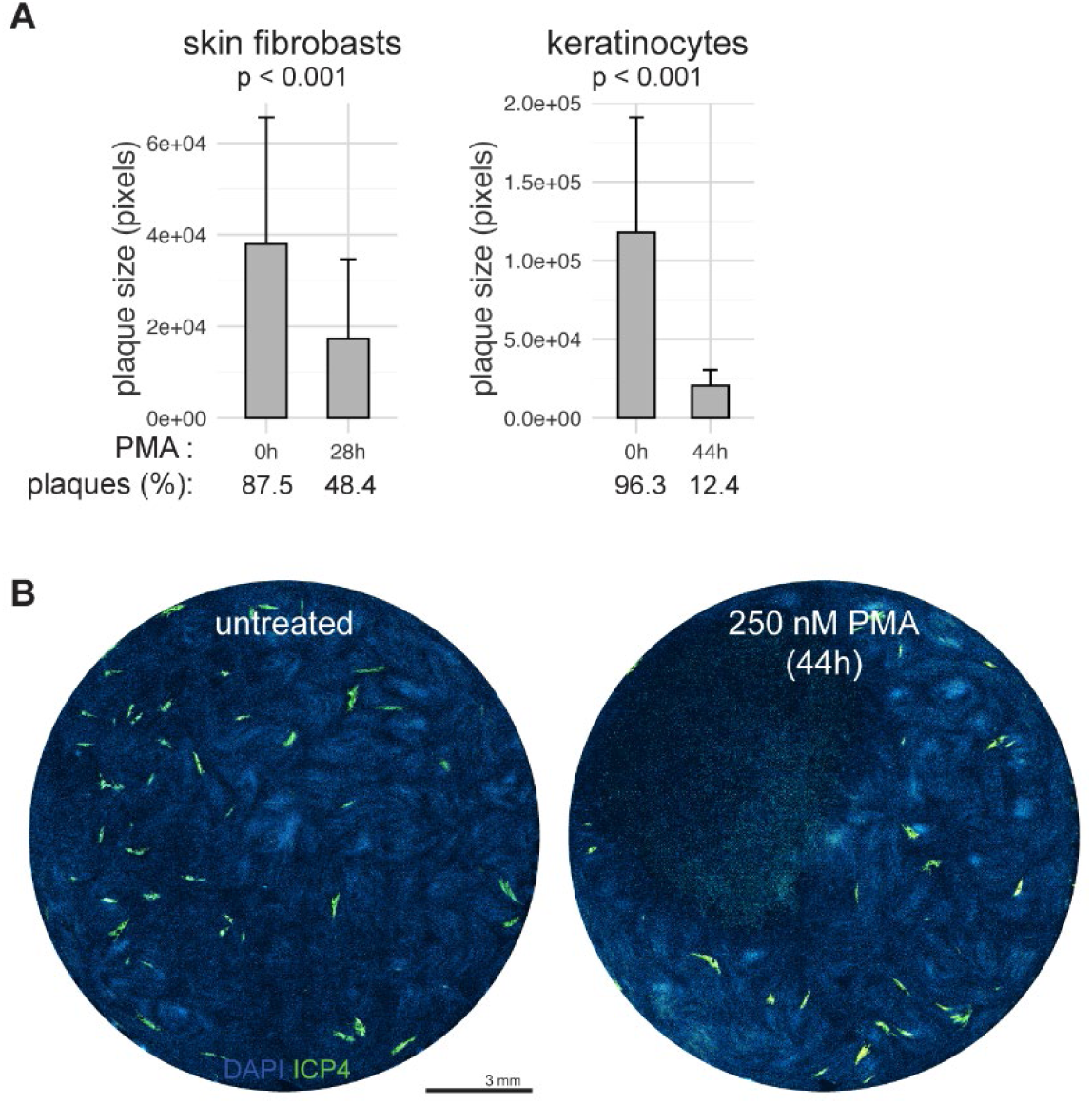
PMA plaque sizes and timing effectiveness. A) : Bar graph of mean + standard deviation of plaque size for PMA-treated and untreated cells. Percentages of plaques are also indicated. B) : Human primary fibroblasts treated with 250 nM PMA for 44 hours in total, 20 hours before cells were infected with WT HSV-1 ICP4-YFP for 24 hours.

**Supplementary Table 1:**
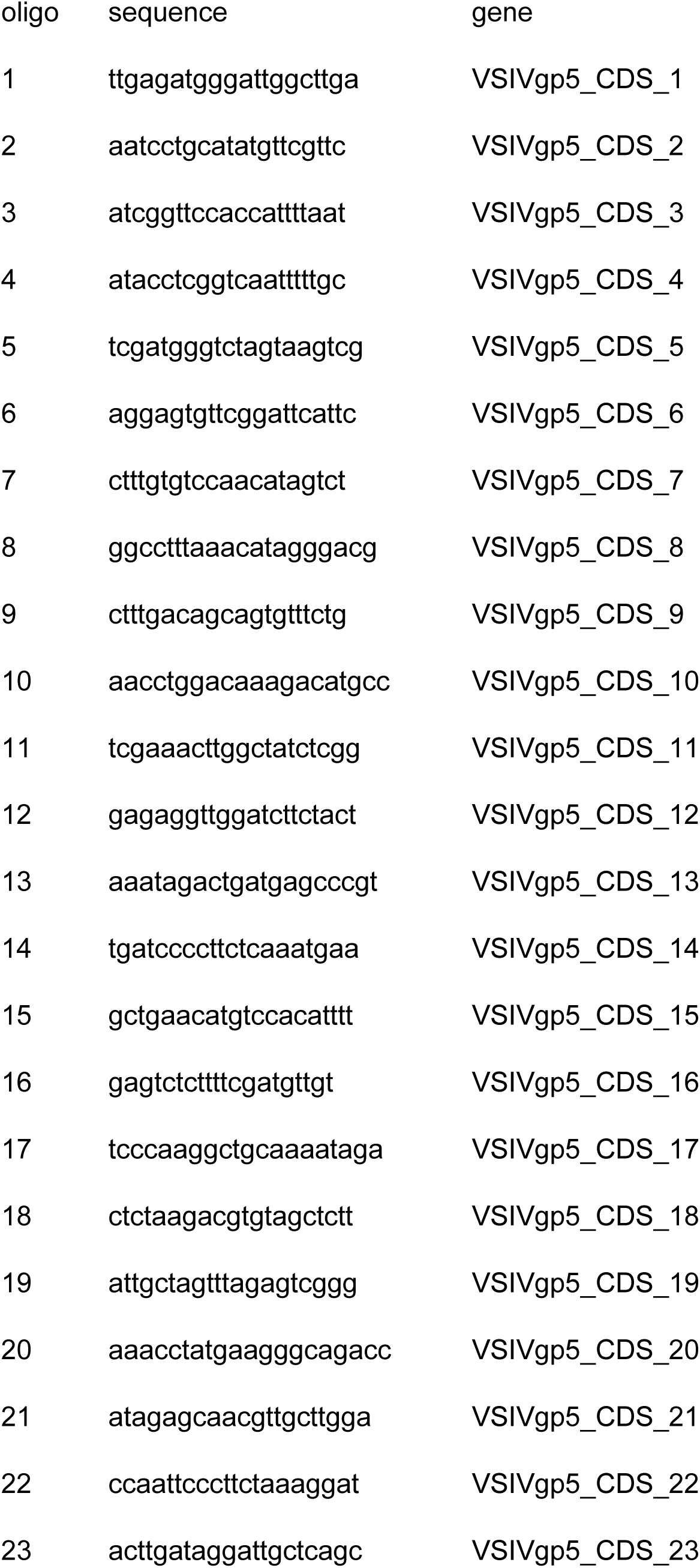
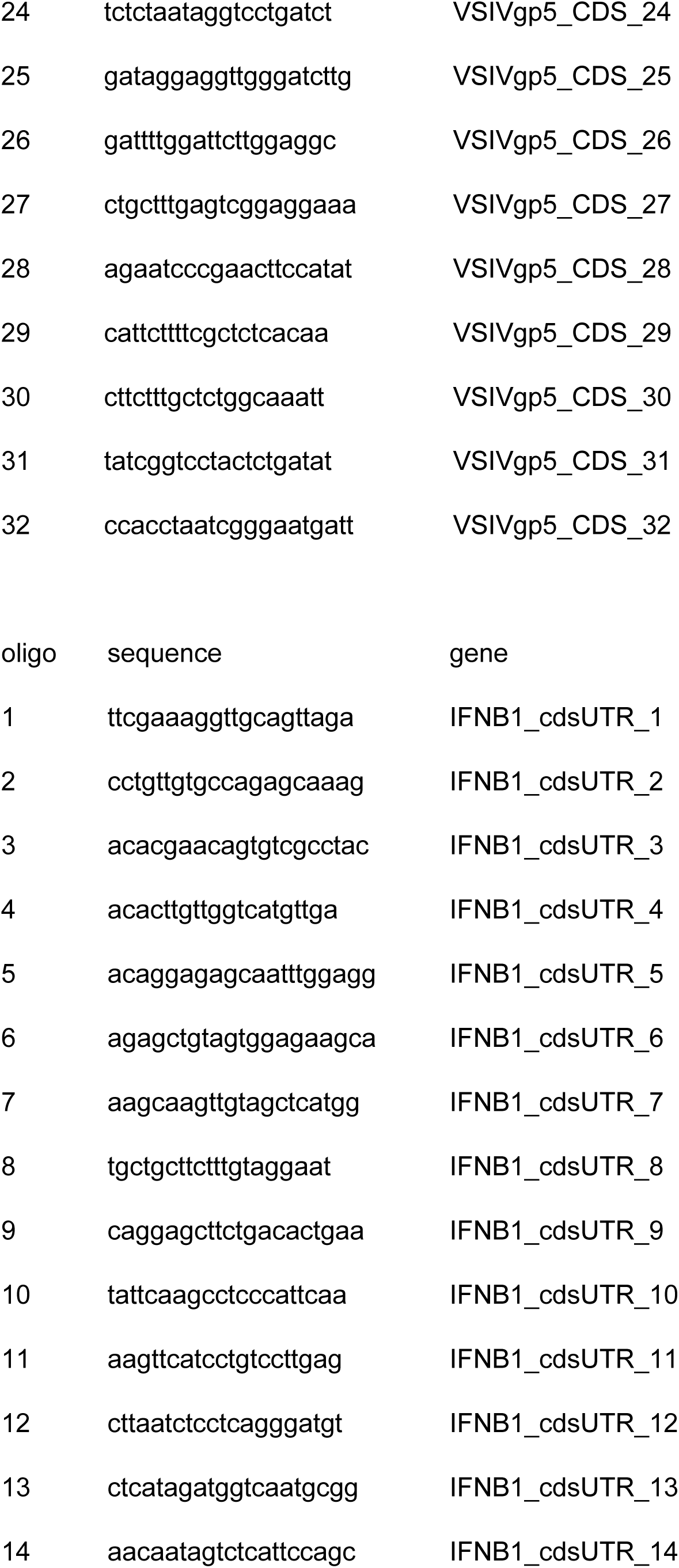
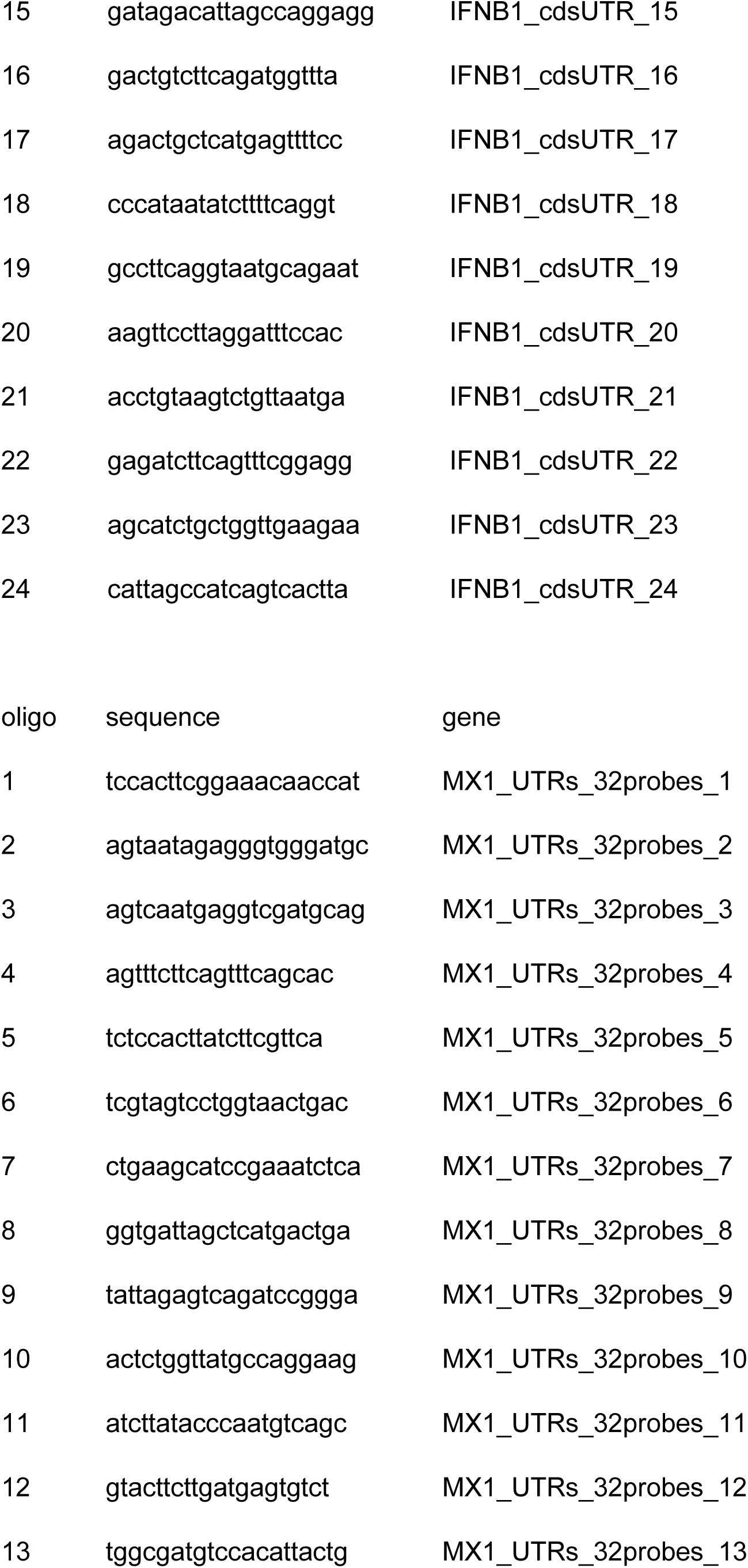
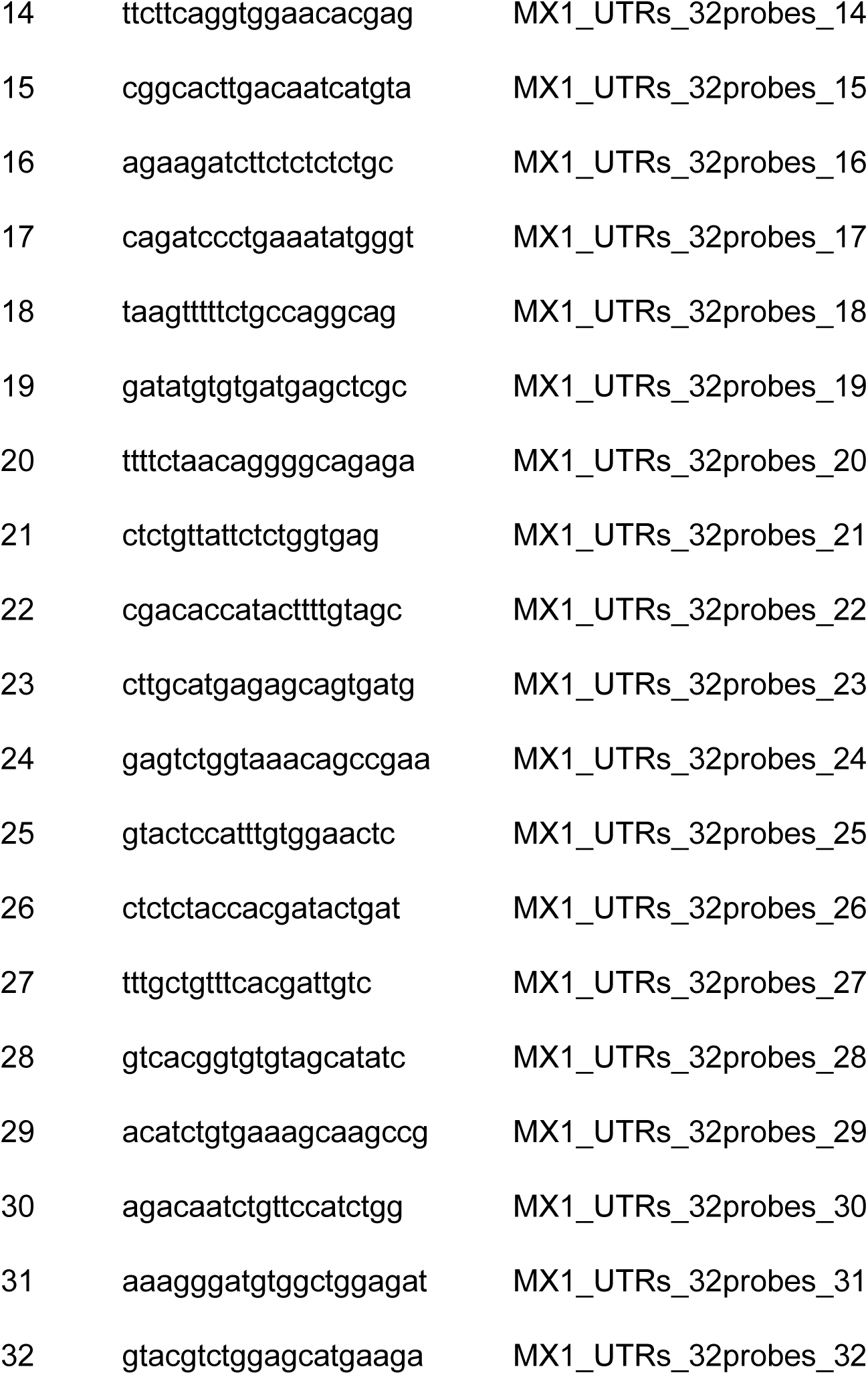
smRNA FISH oligo sequences.

**Supplementary Table 2:**
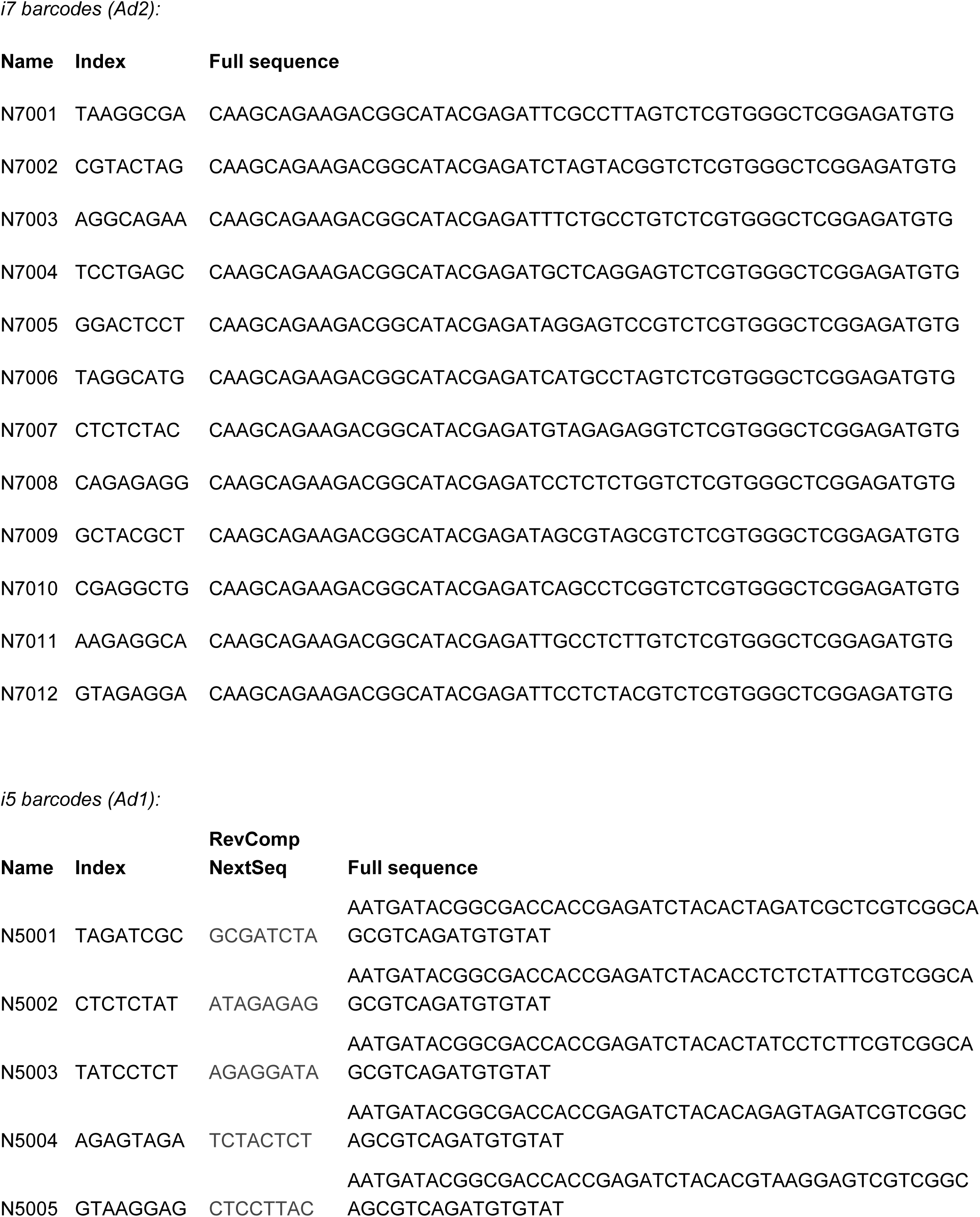
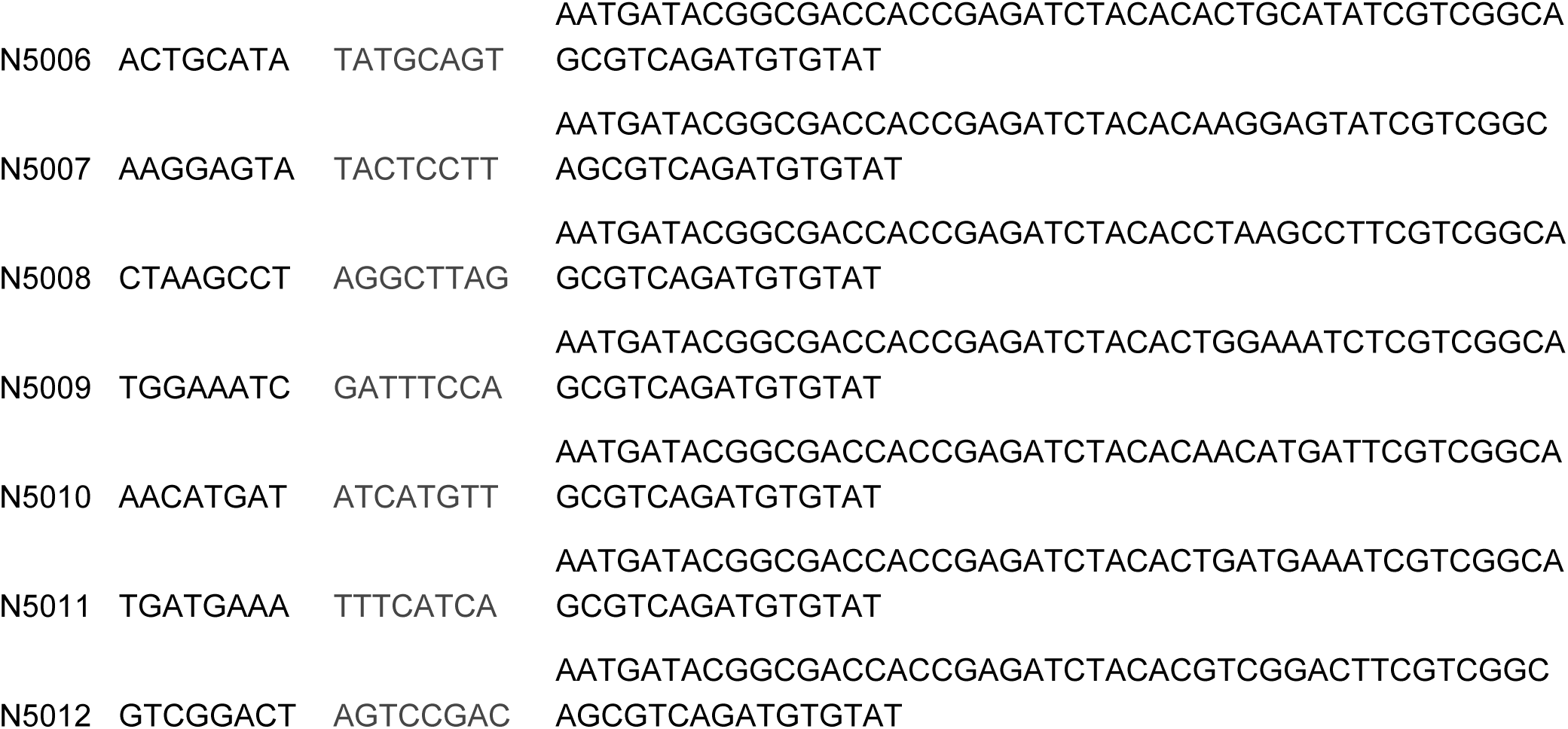
ATAC sequencing primers.

